# Resistance to Spindle Inhibitors in Glioblastoma Depends on STAT3 and Therapy Induced Senescence

**DOI:** 10.1101/2024.06.09.598115

**Authors:** Natanael Zarco, Athanassios Dovas, Virginea de Araujo Farias, Naveen KH Nagaiah, Ashley Haddock, Peter A. Sims, Dolores Hambardzumyan, Christian T. Meyer, Peter Canoll, Steven S. Rosenfeld, Rajappa S. Kenchappa

## Abstract

While mitotic spindle inhibitors specifically kill proliferating tumor cells without the toxicities of microtubule poisons, resistance has limited their clinical utility. Treating glioblastomas with the spindle inhibitors ispinesib, alisertib, or volasertib creates a subpopulation of therapy induced senescent cells that resist these drugs by relying upon the anti-apoptotic and metabolic effects of activated STAT3. Furthermore, these senescent cells expand the repertoire of cells resistant to these drugs by secreting an array of factors, including TGFβ, which induce proliferating cells to exit mitosis and become quiescent—a state that also resists spindle inhibitors. Targeting STAT3 restores sensitivity to each of these drugs by depleting the senescent subpopulation and inducing quiescent cells to enter the mitotic cycle. These results support a therapeutic strategy of targeting STAT3-dependent therapy-induced senescence to enhance the efficacy of spindle inhibitors for the treatment of glioblastoma.

**Highlights:** • Resistance to non-microtubule spindle inhibitors limits their efficacy in glioblastoma and depends on STAT3.

• Resistance goes hand in hand with development of therapy induced senescence (TIS).

• Spindle inhibitor resistant glioblastomas consist of three cell subpopulations—proliferative, quiescent, and TIS—with proliferative cells sensitive and quiescent and TIS cells resistant.

• TIS cells secrete TGFβ, which induces proliferative cells to become quiescent, thereby expanding the population of resistant cells in a spindle inhibitor resistant glioblastoma

• Treatment with a STAT3 inhibitor kills TIS cells and restores sensitivity to spindle inhibitors.

## INTRODUCTION

The prognosis for glioblastoma (GBM) remains dismal (Brown *et al*., 2022), so there is a desperate need to develop more effective therapies. An example is illustrated by a group of anti-proliferative drugs that target the mitotic spindle without affecting microtubules (Liewer and Huddleston 2018; Venere et al., 2015; Gjertsen and Schoffski, 2015), which in this report we refer to as *spindle inhibitors*. These drugs are devoid of the neurotoxicity of microtubule poisons, and include ispinesib, alisertib, and volasertib, which inhibit Kif11, Aurora Kinase A, and Polo Like Kinase, respectively. Although these drugs are CNS penetrant (Gampa *et al*., 2020, Oh *et al*., 2022; Dong *et al*., 2018) and are active against a variety of tumor cells, including GBM stem cells (Dong et al., 2018; Venere et al., 2015; Hong *et al*., 2014), the emergence of treatment resistance has limited their efficacy. We had determined that in the case of the Kif11 inhibitor ispinesib, resistance in GBM depends on two functions of STAT3. The first requires SRC phosphorylation at Y705, which induces STAT3 to stimulate transcription of pro-survival genes. The second requires EGFR pathway-mediated phosphorylation at S727, sending STAT3 to the mitochondria where it activates complexes I and II of the electron transport chain (ETC) and inhibits cytochrome C release in the penultimate stage of apoptosis (Gough *et al*., 2009; Wegrzyn *et al*., 2009; Zhang *et al*., 2013; Kenchappa *et al*., 2022). This explains why combined inhibition of both of SRC and EGFR is required to restore ispinesib sensitivity. The efficacy of ispinesib in orthotopic GBM models can be significantly enhanced by co-administering saracatinib, an CNS penetrant combined SRC/EGFR inhibitor. We also found that ispinesib resistance goes hand in hand with the downregulation of mitotic and spindle checkpoint pathways; upregulation of TGFβ, EMT, and STAT3 pathways; a Proneural-to-Mesenchymal transcriptional shift; resistance to apoptosis; and both cellular and nuclear enlargement.

This work raises new questions that are the focus of our current study. Like ispinesib, both alisertib and volasertib arrest cells in G_2_M, so we wish to know if resistance to these drugs also depends on STAT3. In addition, the cellular enlargement and nuclear atypia that we see in ispinesib resistance are reminiscent of the process of senescence. Although first described as a form of *irreversible* mitotic exit that occurs during aging (Hayflick, 1965), senescence can also develop in malignant cells in response to therapy (Demaria *et al*., 2017; Ruhland and Alspach, 2021), where it is referred to as *therapy induced senescence* (TIS). TIS tumor cells secrete an array of factors, referred to as the *senescence associated secretory phenotype (SASP)*, which supports tumor growth, stemness, and angiogenesis; and which suppresses anti-tumor immunity (Demirci *et al*., 2021; Fitsiou *et al*., 2022; Pribluda *et al*., 2013; Ouchi *et al*., 2016; Salam *et al*., 2023). In addition, one of these SASP factors, TGFβ, induces GBM cells to enter quiescence, a state where cells *reversibly* exit the mitotic cycle (Tejero *et al*., 2019). Since spindle inhibitors are only cytotoxic during the cell cycle, quiescent cells are therefore intrinsically resistant to these drugs. These findings imply that resistance to spindle inhibitors reflects a dynamic interplay between TIS, proliferative, and quiescent subpopulations. We propose that TIS cells resist these drugs by activating STAT3, which suppresses the apoptosis that ordinarily follows prolonged G_2_M arrest. We also propose that through their SASP, TIS cells suppress mitosis in many of the non-TIS cells, further expanding the populations of cells that can resist spindle inhibitors. In this report, we determine that resistance to all three spindle inhibitors relies on both STAT3 and TIS, and have developed a model that may guide future translational applications of these findings.

## RESULTS

### GBM cells resistant to ispinesib, alisertib, and volasertib share a common phenotype

We generated alisertib and volasertib resistant murine *Trp53/Pten*-co-deleted GBM cell lines (*referred to as Trp53/Pten*(−/−)) by exposing naïve cells to drug for three weeks. As in the case of ispinesib, resistance to these other two spindle inhibitors is accompanied by marked cellular and nuclear enlargement and multinucleation in a substantial fraction (**Fig. 1A**), along with increases in pY705 STAT3, pS727 STAT3 (**Fig. 1B, C**), activated SRC, and activated and total EGFR (**Fig**. **1D-J**). Bulk RNAseq of alisertib and volasertib resistant cells reveal patterns of pathway activation very similar to those for ispinesib resistance (Kenchappa *et al*, 2022). These include down-regulation of the mitotic spindle, G_2_M checkpoint, and c-MYC related pathways (*blue arrows in* **Fig. 1K & L**) and up-regulation of STAT3, STAT5, TGFβ, apoptosis, and epithelial-mesenchymal transition (EMT) related pathways (*gold arrows in* **Fig. 1K & L**). Nearly 3000 (53%) of these upregulated genes are shared among spindle inhibitor resistant cells (**Fig. 1M**), consistent with a shared resistance mechanism.

**Figure 1:**
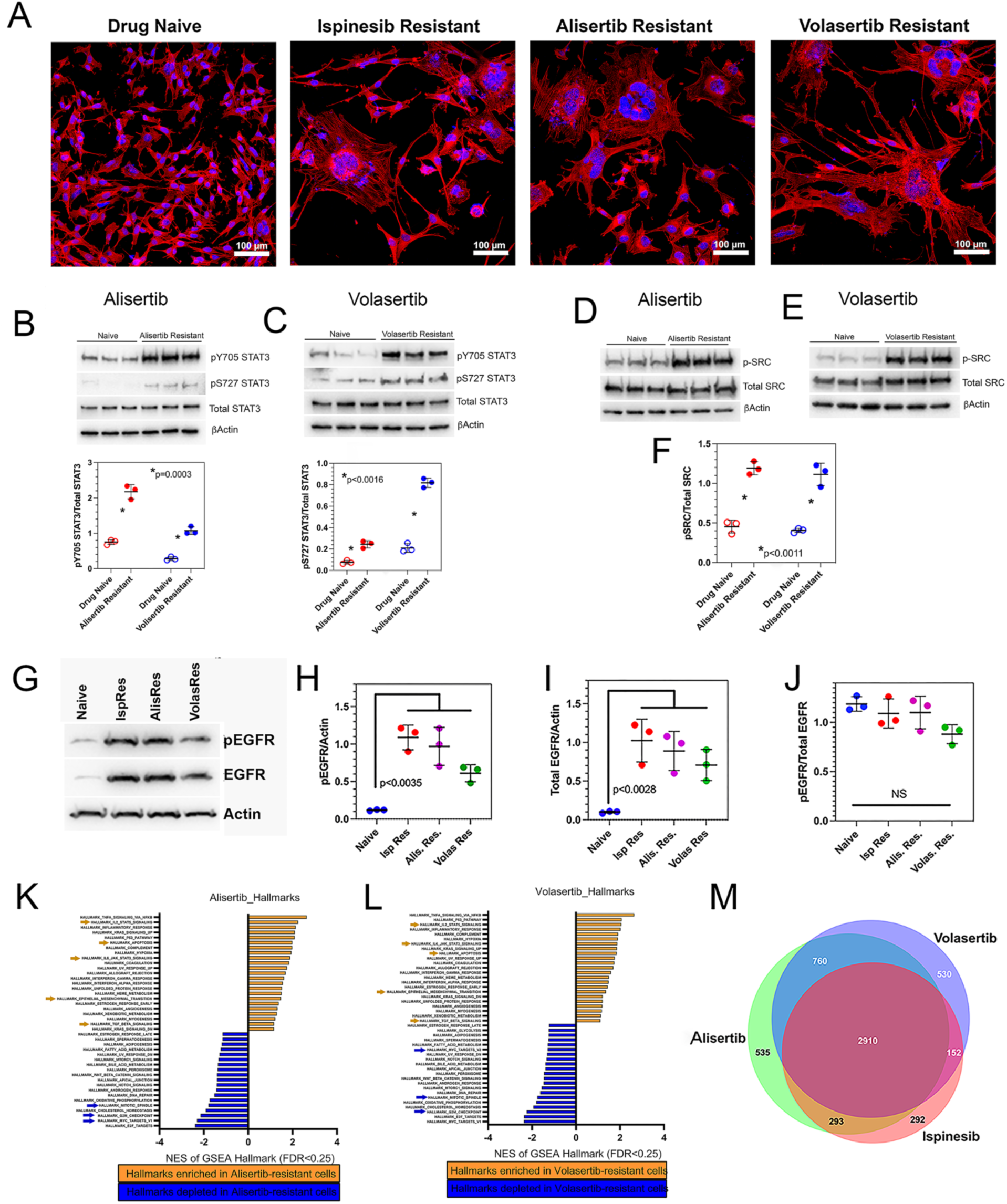
Resistance to three spindle inhibitors works by a similar mechanism. (**A**). Treatment of *Trp53/Pten*(−/−) murine GBM cells with ispinesib, alisertib, or volasertib leads to marked cellular and nuclear enlargement. (**B,C**). Alisertib (**B**) and volasertib (**C**) resistant cells increase STAT3 phosphorylation at Y705 and S727 2-4-fold. (**D-F**). Src phosphorylation is enhanced 2-3-fold in alisertib and volasertib-resistant cells. (**G-J**). The expression of phosphorylated (**G,H**) and total EGFR (**G,I**) increases 4-5-fold in spindle inhibitor resistance. (**K-M**). Gene set enrichment analyses from bulk RNA-seq of alisertib and volasertib-resistant cells (**K,L**). Gold arrows denote upregulated gene ontologies related to STAT3, STAT5, TGFβ, apoptosis, and the epithelial-to-mesenchymal transition, and blue arrows denote downregulated ontologies related to mitotic spindle, G_2_M, and MYC pathways. (**M**). Of those genes upregulated with the development of resistance, nearly 3000 (53%) are shared between ispinesib, alisertib, and volasertib-resistant cells (**M**).

### Resistant GBM cells share therapeutic vulnerabilities in common, reflecting a shared STAT3-dependent mechanism

Ispinesib resistance automatically confers resistance to alisertib and volasertib as well (**Fig. 2A**), which explains why saracatinib (**Fig. 2B**) also reverses alisertib and volasertib resistance. SH5-07, an allosteric inhibitor of STAT3 (Yie *et al*., 2016) that reverses ispinesib resistance (Kenchappa *et al*., 2022) likewise reverses alisertib and volasertib resistance (**Fig. 2C**) in both murine and human (612, L1) GBM lines. Like the murine GBM lines, both of these human lines also upregulate pY705 and pS727 STAT3 upon becoming ispinesib resistant (**Fig. S1**). We also examined if adding saracatinib to alisertib or volasertib improves survival over either drug alone in a *Trp53*-deleted (*Trp53(−/−)*) genetically engineered mouse model (GEMM), as it does with ispinesib (Kenchappa *et al*., 2022). While alisertib (**Fig. 2D**, *left*) alone prolongs median survival over vehicle (*33 versus 45.5 days; p<0.0001 log rank test*), combining it with saracatinib is significantly more effective than alisertib alone (*45.5 versus 53.5 days, p=0.0001, log rank test*). Likewise, combining volasertib (**Fig. 2D**, *right*) with saracatinib significantly prolongs median survival over either drug alone (*volasertib versus saracatinib versus volasertib + saracatinib = 37 vs 34 vs 47 days; p=0.0019, log rank test*).

**Figure 2:**
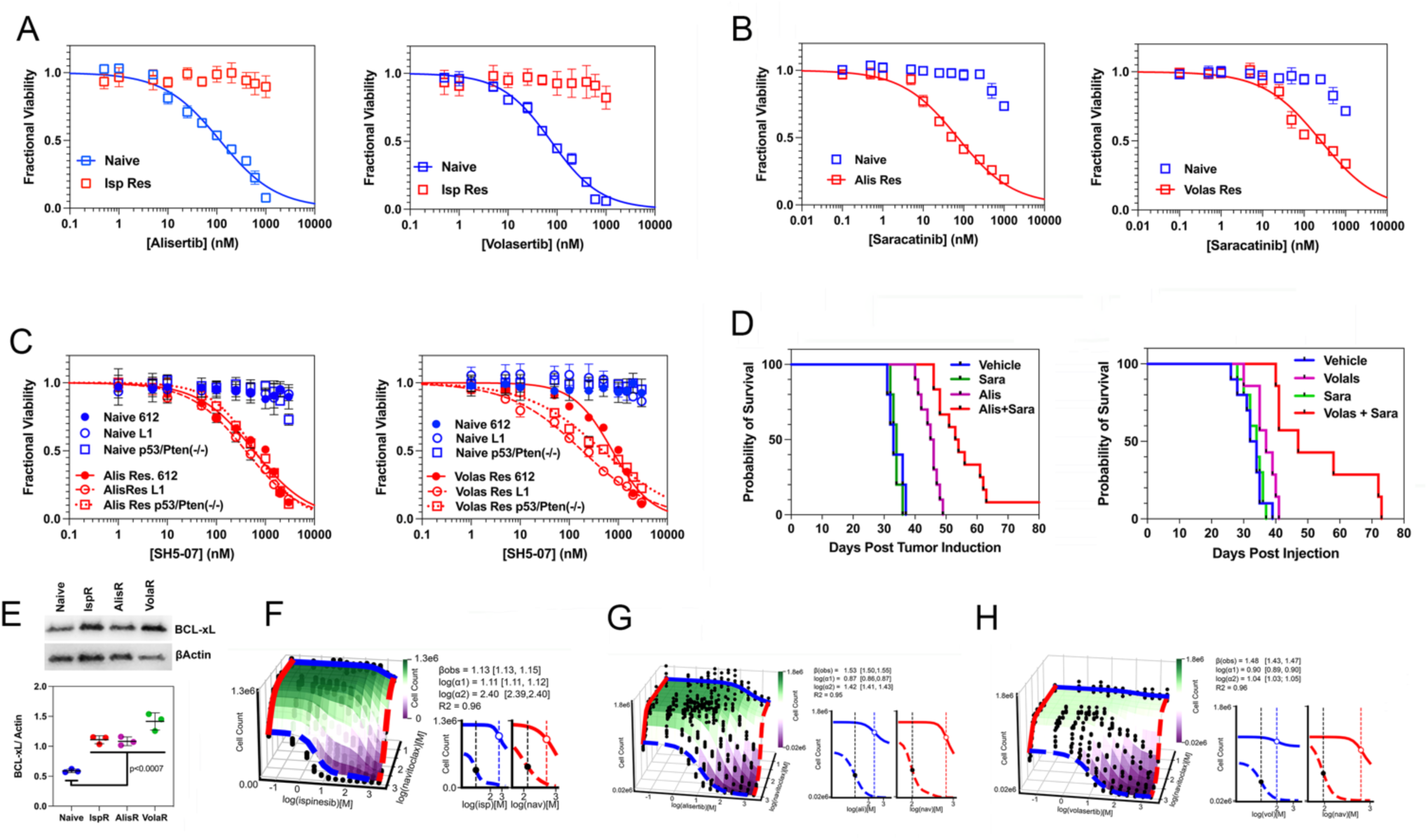
GBM cells resistant to ispinesib, alisertib, and volasertib share therapeutic vulnerabilities. (**A**). Ispinesib-resistant *Trp53/Pten*(−/−) murine GBM cells (*red*) are also resistant to alisertib (*left*) and volasertib (*right*). (**B**). Resistance to alisertib (*left*) and volasertib (*right*) renders cells sensitive to saracatinib. (**C**). While drug naïve human 612 and L1 GBM cells and murine *Trp53/Pten*(−/−) cells (*blue*) are insensitive to the STAT3 inhibitor SH5-07, they become sensitive when resistant to alisertib (*left*) and volasertib (*right*). (**D**). (*Left*). Kaplan Meier survival curves for *Trp53/Pten*(−/−) GEMMs treated with vehicle (*blue*), saracatinib (*green*), alisertib (*magenta*), or alisertib + saracatinib (*red*). (**E**). Spindle inhibitor resistant *Trp53/Pten*(−/−) cells upregulate expression of BCL-xL 2-2.5-fold. (**F-H**). (*Left*) Dose-response surfaces for the combinations of navitoclax with ispinesib (**F**), alisertib (**G**), and volasertib (**H**) in resistant *Trp53/Pten*(−/−) cells. The cell count for each pair of drugs (*black dots*) was fit to the MuSyC equation (*surface plot*), with synergistic combinations denoted by the magenta shading. Values in brackets represent the 95% confidence intervals for the synergy parameter fits.

STAT3 inhibits apoptosis by activating transcription of anti-apoptotic effectors in the nucleus and by preventing opening of the mitochondrial permeability transition pore (MPTP) in the inner mitochondrial membrane (Gough *et al*., 2009; Wegrzyn *et al*., 2009; Zhang *et al*., 2013; Kenchappa *et al*., 2022). Transcription of BCL-xL, one of these anti-apoptotic effectors, is stimulated by pY705 STAT3, and prevents calcium release through the MPTP. This not only blocks cytochrome C release, but it also maintains the mitochondrial membrane redox potential, which drives oxidative phosphorylation (Vander Heiden, *et al*., 1997). BCL-xL can be inhibited by the BCL2 inhibitor navitoclax (Bruncko *et al*., 2007), and its expression is significantly increased in resistant *Trp53/Pten*(−/−) GBM cells (**Fig. 2E**). We therefore examined whether navitoclax reverses resistance by measuring cell viability in the presence of combinations of navitoclax and spindle inhibitors and by fitting data to the synergy algorithm MuSyC (Wooten *et al*., 2021; Meyer *et al*., 2019). MuSyC fits dose-response surfaces (**Fig. 2F-H**, *left*) to drug combination data to calculate the degree of synergistic efficacy (β) and synergistic potency (*log(α12) and log(α21)*). Combining navitoclax with each spindle inhibitor reduces GBM cell viability by >2-fold compared to either drug alone (**Fig. 2F-H**, *right*, *β_obs_ >1.0*) and reduces the EC_50_ of one drug by the other between 7-250-fold (**Fig. 2F-H**, *right*, *log(α12) and log(α21) between. 0.87-2.4*).

### Resistance to spindle inhibitors induces metabolic re-programming

The increase in pS727 STAT3 content with spindle inhibitor resistance predicts that resistance will increase mitochondrial membrane potential, oxidative phosphorylation, and oxygen consumption rate (OCR). Furthermore, EGFR inhibition with erlotinib should reverse these resistance-induced increases by dephosphorylating S727. We measured mitochondrial membrane potential with JC1, a voltage-sensitive mitochondrial fluorophore whose emission shifts from green to red as the mitochondrial inner membrane becomes hyperpolarized. Ispinesib resistance increases the red/green fluorescence ratio nearly 2-fold (**Fig. 3A**). Treatment with 500 nM erlotinib for 24 hours reduces this ratio to that of naïve cells (**Fig. 3B**). While erlotinib has no significant effect on OCR in drug naïve cells (**Fig. 3C**), it reduces maximum OCR by ∼40% in ispinesib resistant cells (**Fig. 3D**, *p=0.011, two-tailed t-test*). Reactive oxygen species (ROS) are a byproduct of oxidative phosphorylation, and we would predict that resistance increases mitochondrial ROS. We stained naïve and ispinesib resistant cells with MitoClox, a mitochondrial localized fluorophore whose emission shifts from red to green with ROS, and found that the green/red intensity ratio increases in resistant cells by 43% (**Fig. 3E**, *p = 0.0003, two-tailed t-test*).

**Figure 3:**
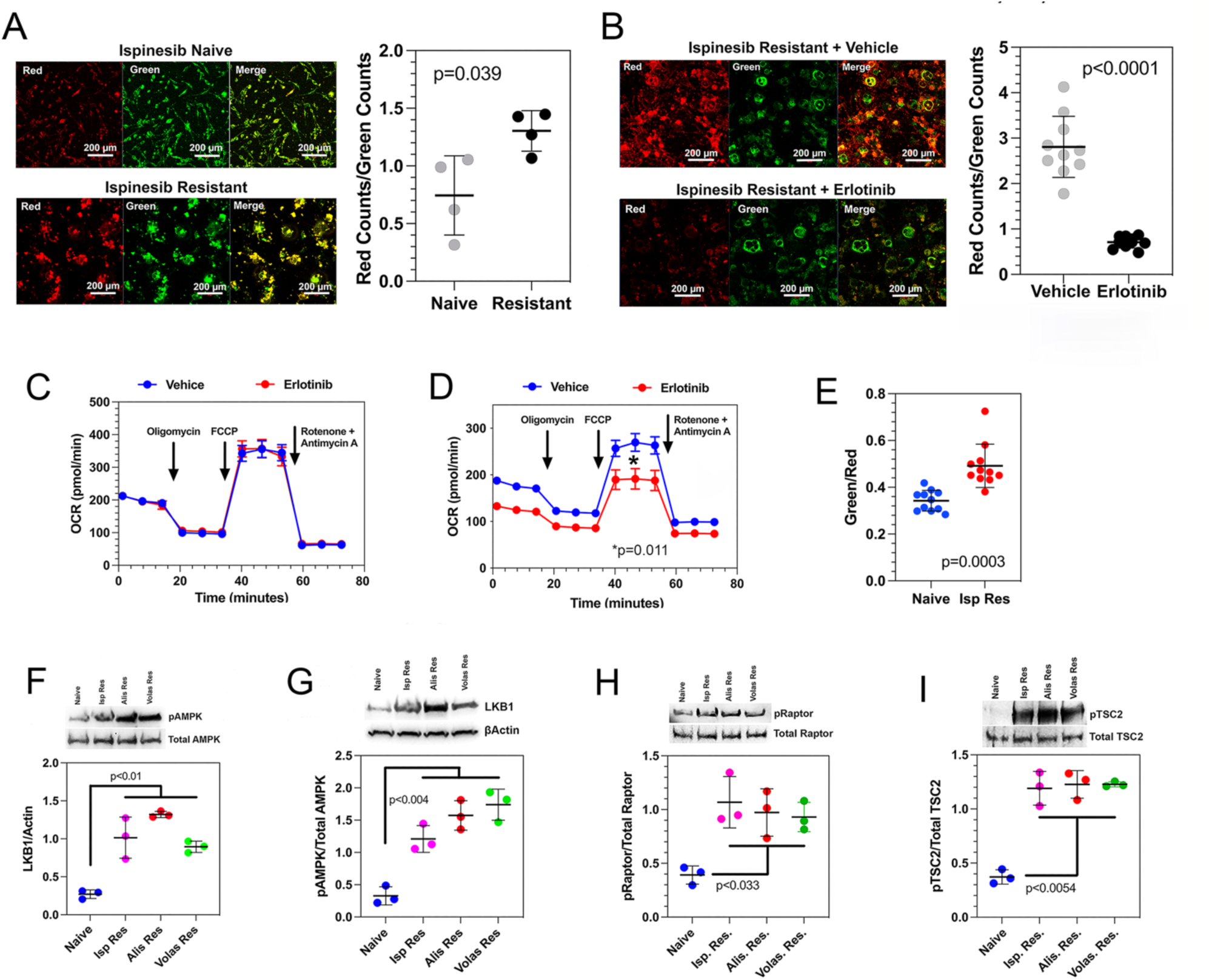
Resistance to spindle inhibitors induces metabolic re-programming. (**A**). JC1 staining of drug naïve (*left, top*) and ispinesib resistant (*left, bottom*) *Trp53/Pten*(−/−) murine GBM cells. Resistance leads to a nearly 2-fold increase in the red to green fluorescence ratio (*right*). (**B**). Compared to vehicle (*left, top*), treatment of ispinesib resistant cells with erlotinib reduces mitochondrial redox potential to levels seen in drug naïve cells (*right*). (**C,D**). Oxygen consumption rate (OCR) for drug naïve *Trp53/Pten*(−/−) murine GBM cells (**C**) and for ispinesib resistant cells (**D**) in the presence of vehicle (*DMSO, blue*) or 500 nM erlotinib (*red*). Error bars represent ± 1 SEM. (**E**). Compared to naïve cells (blue), ispinesib resistance (red) increases mitochondrial ROS, by 43%. (**F-I**). Resistance to spindle inhibitors increases expression of p-AMPK (**F**), p-LKB1(**G**), p-Raptor (**H**), and p-TSC2 (**I**).

GBM cells chronically treated with alisertib reverse the Warburg effect by increasing their reliance on oxidative metabolism for energy production (Nguyen *et al*., 2021). These metabolic changes are accompanied by a reduction in c-MYC, which regulates the Warburg effect (Miller *et al*., 2013). It was proposed that this reflects the loss of active Aurora Kinase A, which binds to c-MYC and protects it from proteasomal degradation. We therefore examined how resistance to ispinesib, alisertib, and volasertib affect levels of c-MYC. In each case, resistance leads to an 8-10-fold reduction in c-MYC expression (**Fig. S2A & B**), consistent with our GSEA analyses (**Fig. 1K & L**). Furthermore, we find that for each inhibitor, resistance leads to a 3-4-fold reduction in Aurora Kinase A expression (**Fig. S2C**).

STAT3 enhances expression of liver kinase B1 (LKB1) and LKB1 activates AMP kinase (AMPK) (Pencik *et al*., 2023). For each spindle inhibitor, resistance leads to a ∼4-fold increase in LKB1 and pAMPK, as well as in two AMPK downstream effectors—TSC2 and Raptor (**Fig. 3F-I**). These findings suggest that STAT3 drives an integrated metabolic response in resistant cells that include enhancement of energy substrate import, through pY705 STAT3 induced transcription of LKB1, and of oxidative phosphorylation through pS727 STAT3 activation of complexes I and II of the ETC.

### Spindle inhibitor resistance and the mesenchymal phenotype reflect a role of STAT3 in both

These results lead us to predict that naïve tumor cells that upregulate pY705 and pS727 STAT3 should *a priori* be more resistant to spindle inhibitors than naïve cells that do not. Furthermore, STAT3 is one of two master transcriptional regulators of the mesenchymal phenotype in GBM (Carro *et al*., 2010), implying that the well-established link between the mesenchymal phenotype and therapy resistance (Pantel *et al*., 2004; Gooding *et al*., 2020; Colman *et al*., 2010) may be mediated by STAT3. While drug naive *Trp53/Pten*(−/−) cells have a Proneural signature, resistance not only to ispinesib (Kenchappa et al, 2022), but also to alisertib and volasertib leads to a Proneural→Mesenchymal shift (**Fig. S3**). We examined the status of STAT3 phosphorylation in four drug naïve GEMM GBM lines. Two of these, MES1861 and MES4622, are Mesenchymal (Gursel et al., 2011; Reilly et al., 2000) and the other two (PN20 and PN24) are Proneural (Hambardzumyan et al., 2009; Chen *et al*., 2023). The content of both pY705 STAT3 and S727 STAT3 is 4-5-fold higher in the MES lines (**Fig. 4A-C**), and this correlates with a 20-50-fold increase in the EC_50_ of the three spindle inhibitors compared to the PN lines (**Fig. 4D-F**; **Table S1**). Suppressing STAT3 with shRNA by >90% (**Fig. S4**) in the two Mesenchymal lines reduces the EC_50_ for each spindle inhibitor 30-50-fold (**Fig. 4G-I**; **Table S1**), to values that are similar to those for the two proneural lines.

**Figure 4:**
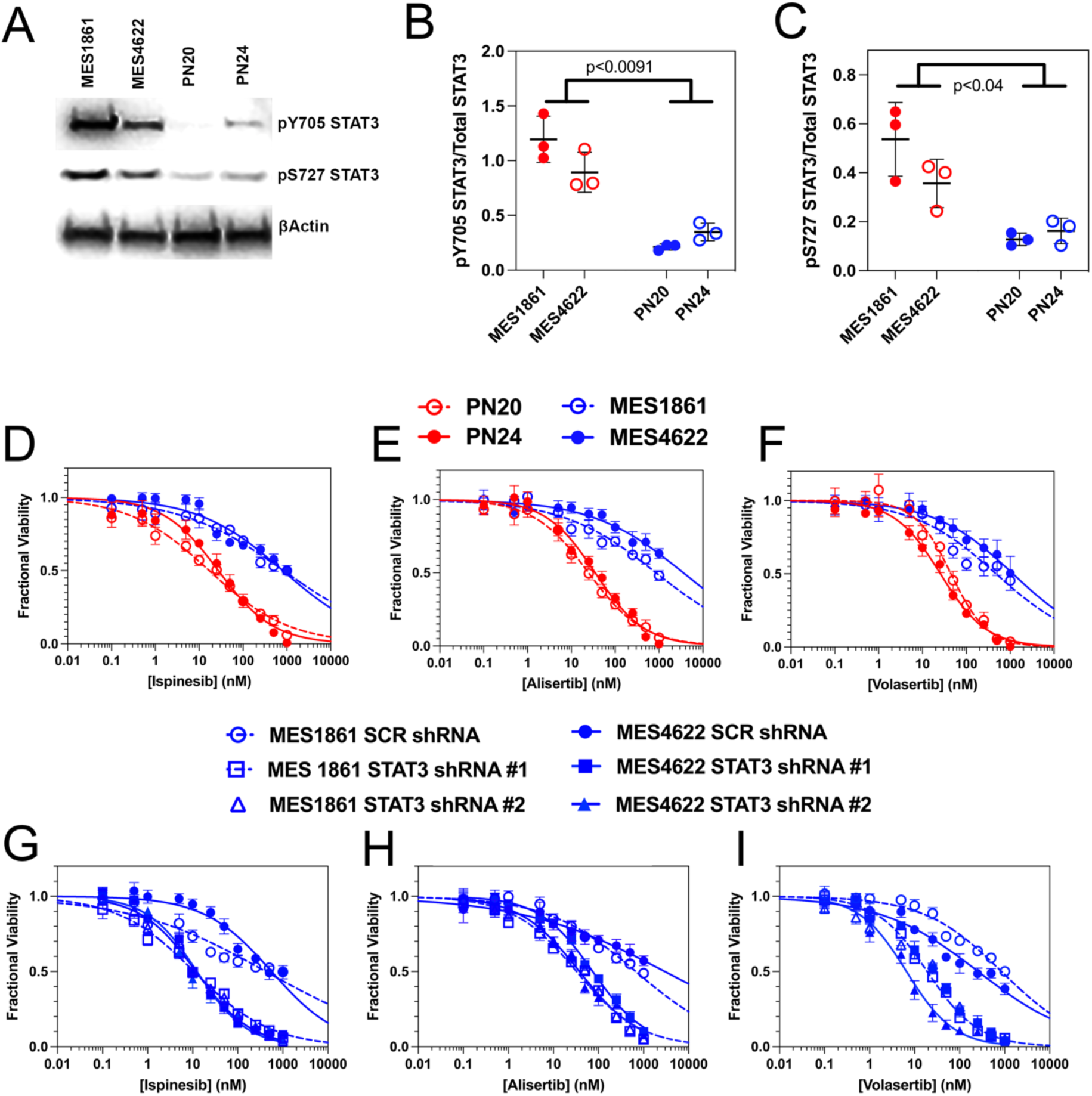
STAT3 activation connects therapy resistance with the mesenchymal phenotype. (**A-C**). Levels of pY705 STAT3 and pS727 STAT3 are increased 4-5-fold in two mesenchymal GEMM lines (MES1861 and MES4622) compared to two proneural GEMM lines (PN20 and PN24). (**D-F**). MES1861 and 4622 lines (*blue*) have EC_50_ values for ispinesib (**D**), alisertib (**E**), and volasertib (**F**) that are 40-50-fold higher than for two proneural lines (*red*). (**G-I**). shRNA suppression of STAT3 in the MES1861 and 4622 lines reduces the EC_50_ for ispinesib (**G**), alisertib (**H**), and volasertib (**I**) by 40-50-fold. See also **Table S1**.

### Resistance to spindle inhibitors is accompanied by therapy induced senescence

The cellular and nuclear enlargement in resistant GBM cells (**Fig. 1A**) has also been described in TIS (Ewald *et al*., 2010). While drug naïve cells are nearly uniformly negative for the senescence marker β-gal (**Fig. S5A***, left*), approximately 50-60% of ispinesib, alisertib, and volasertib resistant cells are positive (**Fig. S5A**, *right;* **Fig. S5B**). Resistance to each inhibitor in *Trp53/Pten*(−/−) GBM cells produces a 4-6-fold increase in the fraction of cells with high forward and side scatter by flow cytometry (**Fig. 5A**), similar to previous reports in TIS (Ewald *et al*., 2010). To determine if these enlarged cells have undergone TIS, we treated both naïve and resistant cells with the fluorescent β gal substrate FDGlu, and performed flow cytometry (**Fig. 5B**, **Fig. S6A**). We plotted the fraction of total cell counts that had high side (**Fig. 5C**) or forward scatter (**Fig. S6B**) and were either β-gal+ (*closed circles*) or β-gal− (*open circles*). We found that nearly all resistant cells with high side and forward scatter are β gal +, while among those with low side (**Fig. 5D**) or forward scatter (**Fig. S6C**) a substantial fraction is also β gal +. This suggests that development of TIS in resistant GBM cells precedes and/or can occur independently of increases in cell size or complexity. We observe similar results a human GBM cell line (L1, **Fig. S7**).

**Figure 5:**
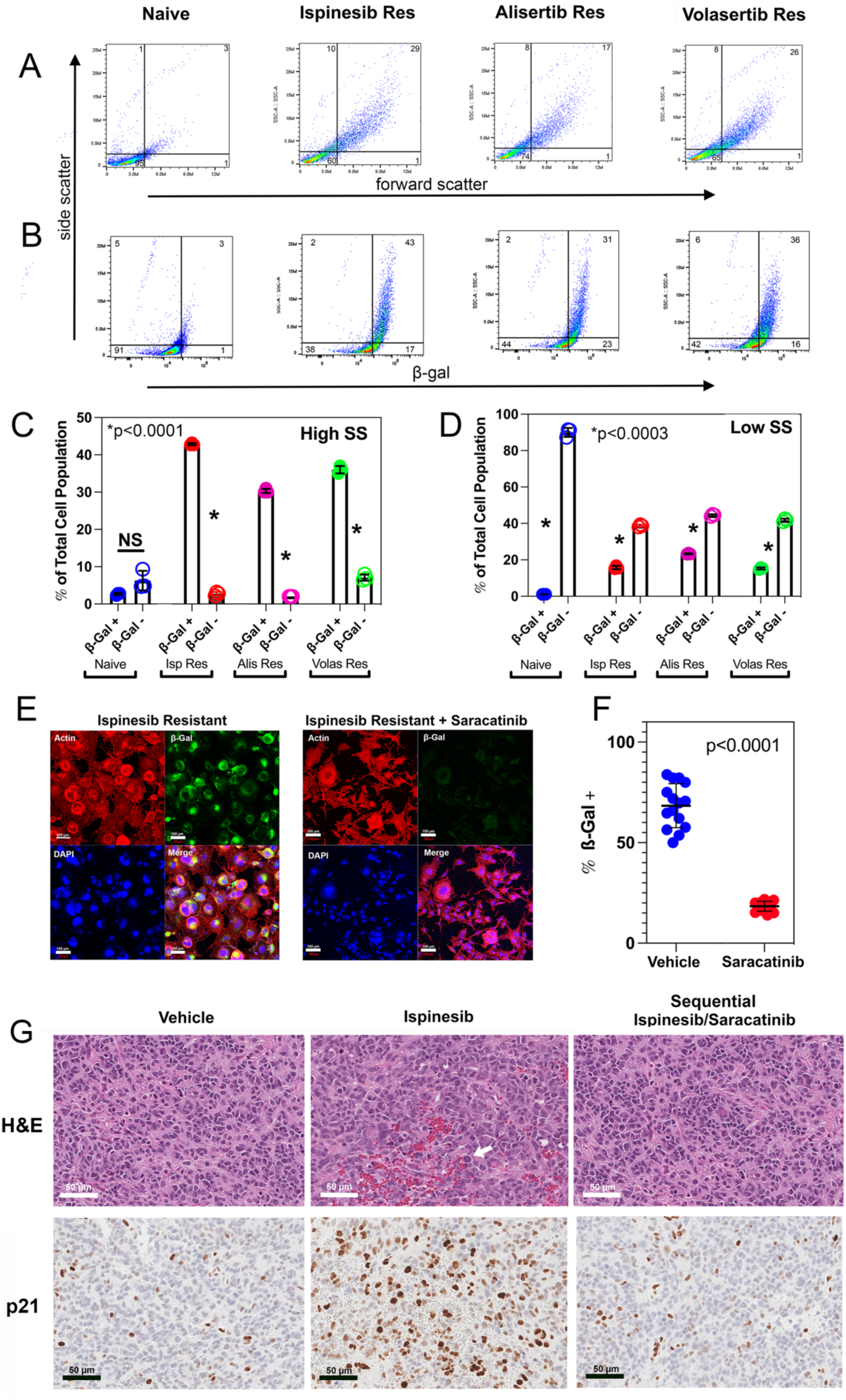
Resistance to spindle inhibitors is accompanied by accumulation of TIS tumor cells. (**A**). Forward *versus* side scatter flow cytometry of naïve and resistant *Trp53/Pten*(−/−). Numbers in each quadrant represent the mean percentages of the total signal. (**B**). Naïve and resistant cells were stained with FDGlu and subjected to flow cytometry to measure side scatter *versus* β-galactosidase activity. (**C**). Bar plot depicting the fraction of the total cell population that demonstrates both high side scatter and either positive or negative β-gal activity. (**D**). Bar plot depicting the fraction of the total cell population that demonstrates both low side scatter and either positive or negative β-gal activity. (**E**). (*Left*) Ispinesib resistant *Trp53*(−/−) cells were stained for actin (*red*), FDGlu (*green*), or DAPI (*blue*). (*Right*) Removal of ispinesib and treatment with 500 nM saracatinib for 72 hours demonstrates near complete loss of β-gal + cells. (**F**). Saracatinib treatment reduces the fraction of cells positive for β-gal activity by > 5-fold. (**G**). GEMMs with orthotopic *Trp53*(−/−) tumors were treated with vehicle, ispinesib x 3 weeks, or with ispinesib x 3 weeks followed by one week of saracatinib. H&E-stained tumor sections reveal that while vehicle treated tumors are composed of cells with relatively uniform size (Fig. 5G*, top left panel*), ∼40% of cells in ispinesib treated tumors have enlarged or multiple nuclei (Fig. 5G*, top center panel*). These cells disappear in tumors treated with ispinesib for three weeks followed by saracatinib for one week (Fig. 5G, *top right panel*). Immunohistochemical staining for p21 (Fig. 5G, *bottom row*) demonstrates sparse staining in vehicle treated mice (Fig. 5G, *bottom left*). Ispinesib markedly increases the number of p21+ cells (Fig. 5G, *bottom center*), while subsequent treatment with saracatinib reduces this number to levels similar to vehicle (Fig. 5G, *bottom right*).

If the TIS subpopulation depends on phosphorylation of STAT3 at Y705 and S727 to prevent apoptosis, then treating resistant tumors with saracatinib should deplete them. We generated ispinesib resistant *Trp53/Pten*(−/−) cells, replaced ispinesib with either vehicle (DMSO) or 500 nM saracatinib for four days, and stained them with DAPI, rhodamine phalloidin, and FDGlu. Results are illustrated in **Fig. 5E & F**. They confirm that saracatinib reduces the fraction of β-gal + cells approximately 5-fold. We treated *Trp*53-deleted GEMMs *in vivo* with vehicle, ispinesib, or with three weeks of ispinesib followed by one week of saracatinib, and examined the brains histologically with H&E and for p21, an immunohistochemical marker of senescence (Althubiti *et al*., 2014). Results are illustrated in **Fig. 5G**. While vehicle treated tumors are relatively uniform in size (**Fig. 5G***, top left panel*), approximately 50% of cells in ispinesib treated tumors contain a substantial fraction of cells with enlarged or multiple nuclei, along with occasional monopolar spindles, a cytological hallmark of ispinesib treatment (**Fig. 5G***, top center panel, white arrow*). These large cells, however, are not apparent in tumors treated sequentially with ispinesib for three weeks followed by saracatinib for one week (**Fig. 5G**, *top*, *right panel*). While vehicle treated tumors show sparse staining for p21 (**Fig. 5G**, *bottom, left panel*), ispinesib increases the p21+ subpopulation significantly (**Fig. 5G***, bottom, center panel*). However, treating with ispinesib for three weeks followed by saracatinib for one week eliminates the vast majority of the p21+ cells (**Fig. 5G***, bottom, right panel*).

### TIS cells are central to the process of spindle inhibitor resistance

Treating ispinesib resistant GBM cells with maximal doses of saracatinib alone eliminates only ∼50% of cells (Kenchappa *et al*, 2022). Eliminating 100% of resistant cells requires combining saracatinib with ispinesib. This suggests that resistant GBMs contain at least two subpopulations—a TIS subpopulation that uses STAT3 to suppress apoptosis and is sensitive to saracatinib, and another that resists ispinesib in some other way and is saracatinib insensitive. Since spindle inhibitors are only active in G_2_M, tumor cells could also resist these drugs by entering quiescence. Quiescence in GBM can be induced by TGFβ and reversed with SB431542, a TGFβ receptor inhibitor (Tejeio *et al*., 2019). The TGFβ signaling pathway is upregulated in GBM cells resistant to ispinesib (Kenchappa *et al*, 2022), alisertib, and volasertib (**Fig. 1K & L**), and TGFβ is a component of the SASP (Pribluda *et al*., 2013; Ewald *et al*., 2010; Fitsiou et al., 2022).

TIS cells can undergo DNA replication by entering the cell cycle at S phase, exiting at G_2_M and re-entering at S phase in a process referred to as *endoduplication*, which leads to nuclear enlargement and multinucleation (Lee *et al*., 2009; Shu et al., 2019). Furthermore, under some circumstances, including loss of *Trp53* (Dirac and Bernard, 2003) or withdrawal of chemotherapy (Duy *et al*., 2021), TIS cells can become proliferative. Regardless, either process can explain why Ki67 can be detected *in vitro* in ispinesib resistant *Trp53*(−/−) GBM cells (**Fig. 6A***, right column*). We also treated orthotopic, *Trp53*(−/−) GEMMs with vehicle (DMSO) or ispinesib for three weeks, and processed the brains for H&E and immunohistochemistry for Ki67. Both vehicle and ispinesib treated tumors stain robustly for Ki67 (**Fig. 6B**), with Ki67 staining seen in ispinesib-treated tumor cells with enlarged nuclei (*black arrows*).

**Figure 6:**
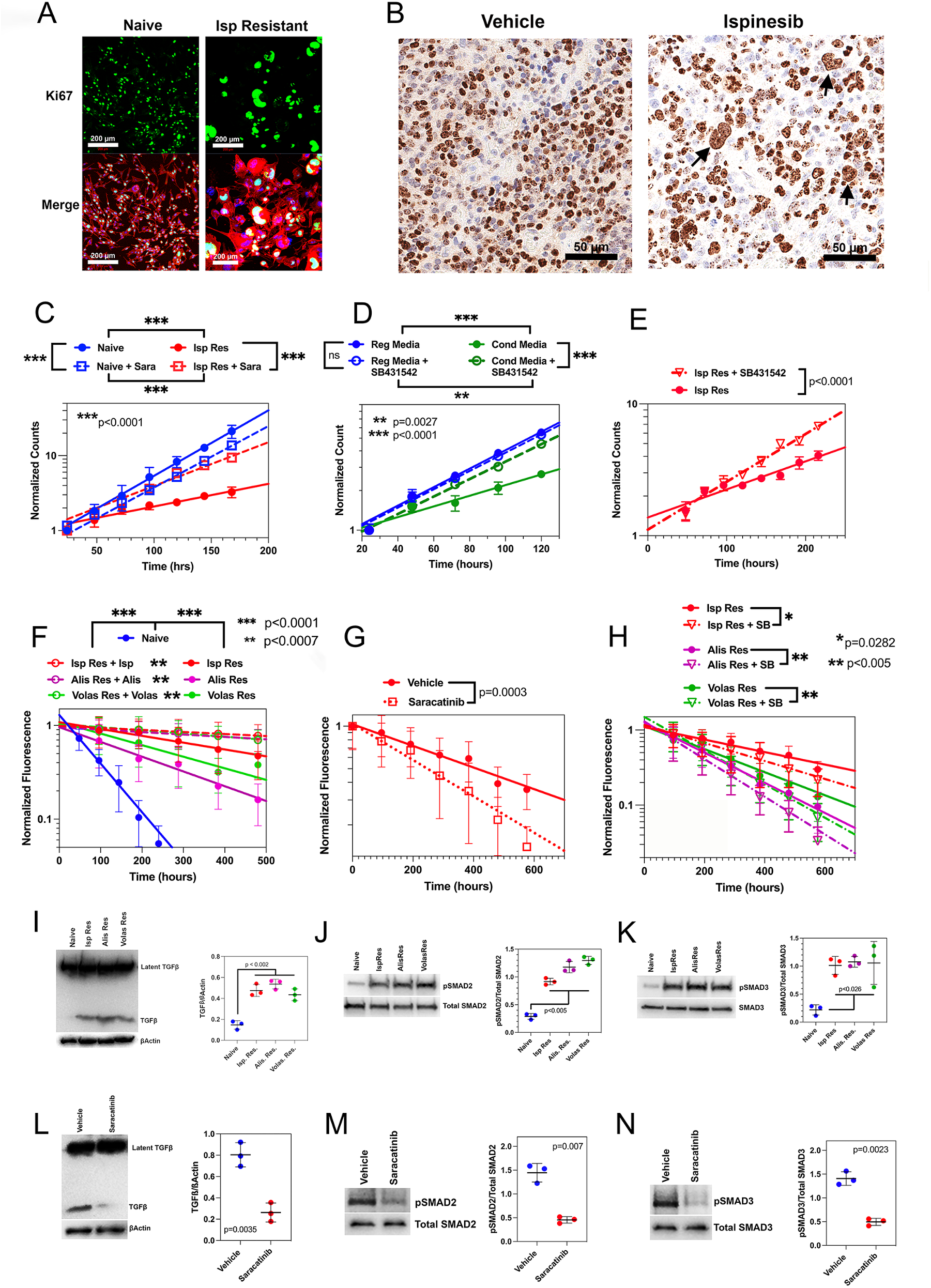
Resistance to spindle inhibitors involves cross talk between SASP components, including TGFβ, and proliferative, TIS, and quiescent tumor cells. (**A**). Ki67 (*top row, green*) staining of drug naïve *Trp53/Pten*(−/−) cells (*left column*) and ispinesib resistant cells (*right column*) *in vitro*. *Bottom row:* merge of Ki67 (*green*) with F actin (*red*) and DAPI (*cyan*). (**B**). Ki67 staining of GBMs from mice treated with vehicle (*left*) or ispinesib (*right*) for three weeks. In the ispinesib treated sample, Ki67 staining can be observed in cells with enlarged (*arrows*) nuclei. (**C-E**). *In vitro* proliferation for *Trp53/Pten*(−/−) cells measured with an ATP-dependent assay (*CellTiter Glo*). (**C**). Proliferation of cells that are naïve (*solid blue circles*), naïve + saracatinib (*open blue boxes*), ispinesib resistant (*solid red circles*), and ispinesib resistant + saracatinib (*open red boxes*), fit to single exponential growth equations (*dashed and solid lines*), (**D**). Proliferation of drug naïve cells in regular media (*solid blue circles*), regular media + SB431542 (*open blue circles*), conditioned media (*solid green circles*), and conditioned media + SB431542 (*open green circles*). (**E**). Ispinesib resistant cells in the absence of ispinesib were treated with vehicle (*solid blue circles*) or SB431542 (*open red triangles*). (**F-H**). Measurement of proliferation using the kinetics of H2B-GFP washout. (**F**). Compared to drug naïve cells (*blue*), cells resistant to ispinesib (*red*), alisertib (*magenta*), or volasertib (*green*) proliferate 2.5-5-fold more slowly in the absence of spindle inhibitors, and ∼16-fold more slowly in their presence. (**G**). Proliferation of ispinesib resistant cells in the absence of ispinesib is accelerated ∼70% by addition of saracatinib. (**H**). Proliferation of ispinesib (*red*), alisertib (*magenta*), and volasertib (*green*) resistant cells in the absence of spindle inhibitor is accelerated 25-45% by addition of SB431542. (**I-K**). Levels of active TGFβ (**I**), phospho SMAD2 (**J**), and phospho SMAD3 (**K**) are increased in cells resistant to ispinesib, alisertib, or volasertib. (**L-N**). In ispinesib resistant cells, treatment with saracatinib significantly reduces active TGFβ (**L**), phospho SMAD2 (**M**), and phospho SMAD3 (**N**).

We measured the proliferation rate of naïve and ispinesib resistant*Trp53/Pten*(−/−) cells in the absence of ispinesib, using an ATP dependent assay (*CellTiter Glo™*) and fit the data to single exponential growth curves. Data are depicted as the solid blue and red lines in the semi-logarithmic plot in **Fig. 6C**. The rate constants for these fits demonstrate that ispinesib resistant cells proliferate ∼4-fold more slowly than naïve cells (**Table S2**). While treating naïve tumor cells with saracatinib reduces proliferation rate by <7% (**Fig. 6C**; *blue open boxes/dashed line*; **Table S2**), it paradoxically accelerates proliferation in ispinesib resistant cells (**Fig. 6C**, *red open boxes/dashed line*; **Table S2**) nearly 2-fold, suggesting that eliminating TIS cells releases mitotic suppression of one or more of the remaining subpopulations.

To test if this suppression of is due to paracrine factors, we compared the proliferation kinetics of drug naïve cells under two conditions—in unconditioned media (*regular media* in **Fig. 6D**) and in media conditioned by exposure to ispinesib resistant cells (*conditioned media* in **Fig. 6D**). Compared to regular media (**Fig. 6D**, *solid blue circles and line*), conditioned media (**Fig. 6D**, *solid green circles and line*, **Table S2**) slows proliferation by >80%. While SB431542 has almost no effect on the proliferation of naïve cells (**Fig. 6D**, *open blue circles, dashed blue line*; **Table S2**), it accelerates proliferation of these cells when cultured in conditioned media by ∼70% (**Fig. 6D**, *open green circles, dashed green line*; **Table S2**). We also find that SB431542 accelerates proliferation of ispinesib resistant cells by ∼70% (**Fig. 6E**, *dashed versus solid red lines*; **Table S2**).

Resistance is associated with increased oxidative metabolism (**Fig. 3**) which leads to increased ATP production, and this could complicate the interpretation of proliferation data using an ATP-dependent assay. We therefore also used a second method to measure proliferation, which involves transfection of drug naive *Trp53/Pten*(−/−) murine or L1 human GBM cells with a lentiviral vector encoding an H2B-GFP fusion protein under transcriptional control of the doxycycline promoter. Briefly treating with doxycycline to induce H2B-GFP expression and then removing it leads to loss of GFP fluorescence over time due to its serial dilution in generations of proliferating daughter cells. An example is depicted in **Fig. S8A** for drug naïve and resistant L1 cells. Fluorescence intensity data were fit to single exponential decays to yield apparent rate constants for proliferation, summarized in **Table S2** for *Trp53/Pten*(−/−) murine and in **Table S3** for L1 human GBM cells. Although there are differences between corresponding rates for the ATP-dependent and H2B-GFP pulse chase methods, both show that the proliferation of resistant cells in the absence of spindle inhibitor is ∼3-4-fold slower than for naïve cells (**Fig. 6F**, **Fig. S8B**), and that both saracatinib (**Fig. 6G**) and SB431542 (**Fig. 6H**, **Fig. S8C**) accelerate the proliferation of resistant cells. Furthermore, the proliferation of resistant cells in the presence of a spindle inhibitor is 5-15-fold slower than in its absence (**Fig. 6F**, **Fig. S8B**, *dashed versus solid lines*).

These results imply that TIS cells can suppress proliferation in the non-TIS population through paracrine factors. In support of this, levels of active TGFβ as well as of two of its downstream effectors, SMAD2 and SMAD3, are 4-5-fold higher in resistant *Trp53/Pten*(−/−) cells cultured without spindle inhibitors than in drug naïve cells (**Fig. 6I-K**). Treatment with saracatinib reverses these effects (**Fig. 6L-N**).

### The interactions between proliferative, quiescent, and TIS GBM cells support a sequential treatment strategy

Our results support the model depicted in **Fig. 7A**. We propose that GBMs consist of proliferative, quiescent, and TIS subpopulations. While some proliferative cells are killed by spindle inhibitors, others activate STAT3 to suppress the apoptosis that typically follows a prolonged G_2_M block and enter a TIS state. TIS cells in turn suppress the conversion of quiescent to proliferative cells through components of the SASP, including TGFβ. We also propose that proliferative cells are in a reversible equilibrium with quiescent cells. Whether the TIS state can revert to the proliferative (*indicated by the thin arrow in* **Fig. 7A**) or not, our model predicts that depleting proliferative cells with a spindle inhibitor will reduce the fraction of this subpopulation and increase the fraction of TIS cells. Conversely, killing TIS cells with saracatinib should reverse the TIS-induced block on the quiescent→proliferative transition, leading to a corresponding increase in the proliferative compartment.

**Figure 7:**
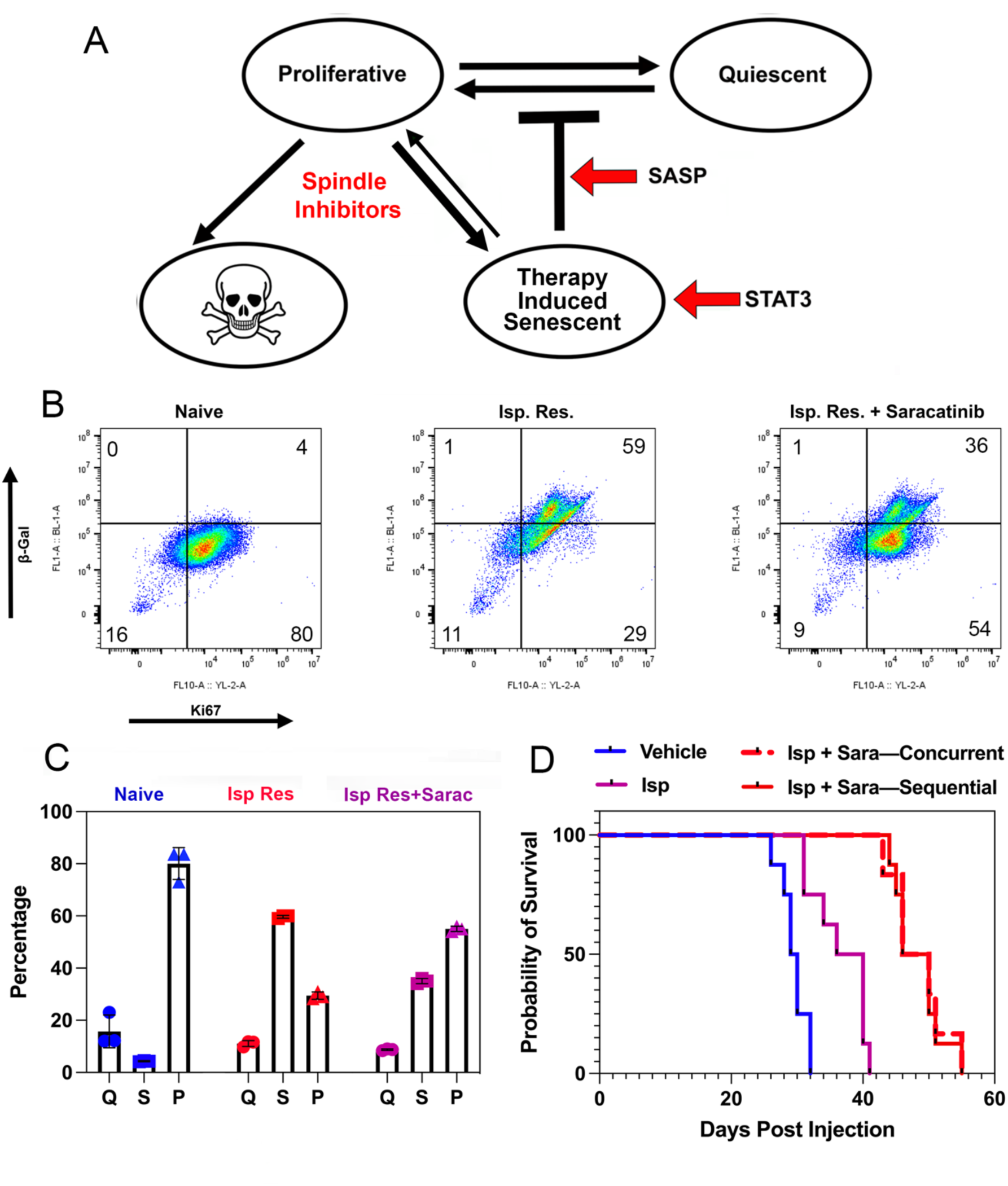
The interactions between proliferative, quiescent, and TIS GBM cells support a sequential treatment strategy. (**A**). Three cell state model for resistance to spindle inhibitors. (**B**). Two color flow cytometry scatter plot of naïve, ispinesib resistant, and ispinesib resistant +saracatinib treated *Trp53*(−/−) cells. Numbers in each quadrant represent the mean percentages of the total signal. (**C**). Percentages of the total cell population that are quiescent (*left*), senescent (*center*), or proliferative (*right*), for naïve (*blue*), ispinesib resistant (*red*), and ispinesib resistant treated with saracatinib for 48 hours (*magenta*). (**D**). GEMMs with orthotopic *Trp53*(−/−) GBMs were treated with vehicle (*blue*), ispinesib (*magenta*), ispinesib + saracatinib given concurrently (*red dashed*), or in alternating cycles of 3 weeks of ispinesib followed by 1 week of saracatinib (*solid red*).

We have tested these predictions by performing flow cytometry on three groups of *Trp53/Pten*(−/−) cells—naïve, ispinesib resistant, and ispinesib resistant treated with saracatinib for 48 hours. Cells were stained with FDGlu, to mark β-gal + cells, and anti-Ki67, to mark cycling cells, and results are depicted in **Fig. 7B**. The four quadrants in the two-dimensional scattergram correspond to the following: <u>lower left:</u> Ki67− and β-gal− (*quiescent*); <u>lower right:</u> Ki67+ and β-gal− (*proliferative*); <u>upper left:</u> Ki67− and β-gal+ (*TIS*); and <u>upper right:</u> Ki67+ and β-gal+ (*TIS cells undergoing DNA replication*). **Fig. 7C** illustrates the percentage of TIS *(β-gal +, Ki67 + and −*), proliferative, and quiescent subpopulations for naïve (*blue*), ispinesib resistant (*red*) and ispinesib resistant + saracatinib (*magenta*) tumor cells. Ispinesib resistance reduces the fraction of proliferative cells ∼3-fold (*p=0.0017, two tailed t test*) and increases the fraction of TIS cells ∼10-fold (*p<0.0001, two tailed t test*), consistent with some of the former evolving into the latter. Treating ispinesib resistant cells with a brief exposure to saracatinib (*48 hours*) partially reverses these effects.

Our model explains why saracatinib paradoxically stimulates proliferation in resistant cells but not in drug naïve cells (**Fig. 6G, K**), since reducing the TIS subpopulation with saracatinib would reduce SASP factors and release quiescent cells to proliferate. Our model also explains the survival benefit of combining spindle inhibitors with saracatinib (**Fig. 2G & H** & Kenchappa *et al*, 2022), which would deplete the TIS subpopulation, enabling more quiescent cells to become proliferative and become spindle inhibitor sensitive. One problem with translating this approach clinically is that both spindle inhibitors and the senolytics saracatinib and navitoclax have overlapping toxicities, including myelosuppression (Blagden *et al*., 2008; Kaye *et al*., 2012; Mosse *et al*., 2019; Wilson *et al*., 2010). However, administering the same doses of these two drugs in an alternating schedule might not only provide equivalent survival benefit with less toxicity, but also allow dose escalation of each drug beyond what could be tolerated when both are co-administered. As a first step to test this, we treated our *Trp53*(−/−) GEMMs with vehicle, ispinesib alone, concurrent ispinesib + saracatinib, or with repeated cycles of ispinesib for three weeks alternating with saracatinib for one week. The resulting Kaplan Meier curves (**Fig. 7D**) show that the concurrent and sequential treatment schemes produce statistically indistinguishable survival benefits (*median survival 48 days, p=0.82, log rank test*), which are both significantly better than ispinesib or vehicle alone (*p<0.0005, log rank test*).

## DISCUSSION

### Spindle inhibitor resistance works by a shared set of mechanisms and generates a shared set of vulnerabilities

Although ispinesib, alisertib, and volasertib inhibit distinct components of the mitotic spindle, resistance to each produces the same phenotype, including upregulation of nearly 3000 genes in common as well as cytomegaly and nuclear pleomorphism (**Fig. 1**). This argues that the mechanism of resistance to these drugs does not involve mutations that prevent drug binding to target and subsequent blocking of mitotic progression, a point we established previously with ispinesib resistance (Kenchappa *et al*., 2022). Rather, our data suggests that resistance works by blocking the apoptosis that ordinarily follows a prolonged G_2_M arrest. Furthermore, our finding that ispinesib resistance confers resistance to both alisertib and volasertib as well indicates that resistance to each inhibitor relies upon the same mechanism.

The gene sets enriched in resistant cells include those involved in the IL6-JAK-STAT3 pathway, which our prior scRNA-seq studies (Kenchappa *et al*., 2022) found are uniformly upregulated in ispinesib-resistant murine *Trp53/Pten*(−/−) GBM cells, as well as those involved in apoptosis pathways. Among the latter is the anti-apoptotic protein BCL-xL, which is transcriptionally regulated by STAT3 (Catlett-Falcone, *et al*., 1999). Our finding that the BCL-xL inhibitor navitoclax is highly synergistic with each spindle inhibitor supports our proposal that increasing the transcription of anti-apoptotic effectors is necessary for resistant GBM cells to survive. This conclusion is also supported by our results which show that STAT3 inhibition with saracatinib or SH5-07 is specifically cytotoxic to resistant cells.

### STAT3 activation explains the association of treatment resistance with the mesenchymal phenotype and metabolic reprogramming

An additional set of genes that are upregulated in resistant cells are those related to the epithelial-mesenchymal transition (EMT). EMT is frequently associated with development of resistance to chemotherapy and radiotherapy (Brabletz *et al*., 2018), and a similar Proneural→Mesenchymal shift has been associated with treatment resistance in GBM (Kim *et al*., 2021). STAT3 and CEBPβ are the two master regulators of the Mesenchymal phenotype in GBM (Carro, *et al*., 2010), and as we have shown (**Fig. 1B-E** and Kenchappa, *et al*., 2022), activation of STAT3 and expression of the IL6-JAK-STAT3 pathway occurs with spindle inhibitor resistance. In addition, our bulk RNA-seq data demonstrate that CEBPβ is also upregulated in ispinesib, alisertib, and volasertib resistant cells (*Gene Expression Omnibus sequential accession numbers GSM8267005-GSM8267016*). We thus propose that activation of STAT3 underlies the association between resistance and the mesenchymal phenotype (**Fig. 2E-H**). While this suggests that mesenchymal GBMs are resistant to spindle inhibitors because of STAT3, it does not necessarily follow that all GBMs resistant to spindle inhibitors are mesenchymal. This is highlighted by our recent study (Cheng *et al*., 2024) which showed that ispinesib resistance in TS543, a proneural human GBM line, does not lead to a Proneural→Mesenchymal shift. BCL-xL expression can be enhanced not only by STAT3, but also by other transcription factors, including MYB, which is upregulated in ispinesib-resistant TS543 cells (Yuan *et al*., 2010). This suggests that the final common pathway for spindle inhibitor resistance may involve anti-apoptotic proteins, including BCL-xL, whose expression is STAT3-dependent in tumors that upregulate STAT3, and STAT3 independent in some non-mesenchymal tumors.

Resistance induces metabolic changes designed to enhance energy production, including an increase in activated AMPK. Furthermore, resistance to both ispinesib (**Fig. 3A-E**) and alisertib (Nguyen *et al*., 2021) increases oxidative metabolism, which in the case of alisertib is driven by fatty acid oxidation. As we have shown, resistance to each of these spindle inhibitors also leads to TIS, and TIS in turn enhances lipid oxidation (Flor *et al*., 2017). Resistance leads to a downregulation of G_2_M checkpoint and mitotic effectors (**Fig. 1K & L**; Kenchappa et al., 2022) including Aurora Kinase A. If Aurora Kinase A protects c-MYC from proteasomal degradation (Nguyen et al., 2021), then its downregulation with resistance (**Fig. S2C**) could explain the loss of c-MYC that we observe, leading to a reversal of the Warburg effect and a shift toward a more oxidative metabolic profile.

### Therapy induced senescence accompanies the development of resistance

Like normal cells that undergo senescence during aging, tumor cells that undergo TIS express senescence markers, such as βgalactosidase and p21. Furthermore, like their normal counterparts, TIS cells are deleterious, as they produce immunosuppressive, angiogenic, and stemness supporting effectors as part of the SASP (Ruhland and Alspach, 2021). However, analogies between normal and malignant cells only go so far. For example, while p53 is needed for senescence in aging, it is not for malignant cells in a TIS state. Likewise, as noted above, TIS cells can remain Ki67+ by replicating their DNA through endoreduplication or by reverting to a proliferative phenotype (Saleh *et al*., 2019, 2020; Roberson *et al*., 2005; Alotaibi *et al*., 2016; Shu *et al*., 2018; Holland and Cleveland, 2012). As with STAT3 and the Mesenchymal phenotype, the development of TIS on the one hand and of large size and complexity on the other may not be strictly linked, since flow cytometry shows that development of TIS can precede these morphologic changes (**Fig. 5 A-D, Figs. S5 & S6**).

### Resistance to spindle inhibitors in GBM can be described by a three-cell state model that has translational ramifications

GBMs consist of dynamic cellular subpopulations, and this has led to classification schemes that sort GBM cells into subgroups based on their lineage resemblance or functionality (Patel *et al*., 2014; Neftel *et al*., 2019; Yuan *et al*., 2018; Garofano *et al*., 2021). Nevertheless, one feature common to all of these is proliferation. For example, one scheme (Neftel *et al*., 2019) described four subgroups, two of which (*Astrocyte-like and Mesenchymal-like*) are considerably less proliferative than the other two (*NPC-like and OPC-like*). Likewise, one subgroup in another classification scheme (Garofano *et al*., 2021) is distinguished from the others by its strong proliferative signature. Thus, sorting GBM cells into proliferative and or non-proliferative subgroups is consistent with each of these schemes. In our model of resistance, we further subdivide the non-proliferative subgroups into quiescent and TIS, which should both resist spindle inhibitors. These considerations lead to the three-cell state model in **Fig. 7A**, whose salient features are supported by both our *in vivo* (**Fig. 6 A & B**) and *in vitro* (**Fig. 7 B & C**) data.

The effect of resistance on cell growth is relatively durable, since proliferation rates after removal of spindle inhibitors are ∼4-fold slower than for naïve cells (**Fig. 6C & F**, **Fig. S8B**). However, in the presence of spindle inhibitors, these rates become ∼15-20-fold slower than drug naïve cells (**Tables S2 & S3**). We propose that these growth rates represent an ensemble average of three intrinsic rates: the growth rate of proliferative cells and the rates of conversion of quiescent and TIS cells to proliferative cells. In the presence of spindle inhibitors, the population of proliferative cells drop, since some are killed by the drug and others develop TIS. In this case, cell growth rate would be largely determined by the transition rate from the TIS to the proliferative state. If so, our results would suggest that this would be an uncommon event, accounting for the overall very slow proliferation rate that we observe (**Tables S2 & S3***; thin arrow in* **Fig. 7A**). With the removal of spindle inhibitor, any cells remaining in the proliferative compartment could grow unimpeded. However, the number of cells in this compartment would be appreciably smaller than for naïve cells, due to their depletion by prior exposure to spindle inhibitor (**Fig. 7C**); and this would reduce the value of the overall, ensemble averaged growth rate. Furthermore, conversion of quiescent cells to proliferative would still be blocked by the presence of TIS cells. This would result in an overall growth rate that is still slower than the original, drug naïve population.

Given its effects in eliminating TIS cells, saracatinib could be considered a “senolytic”. However, not all senolytics are equally effective in all contexts. This is highlighted by our prior study (Kenchappa et al., 2022) which showed that dasatinib, a SRC family kinase inhibitor which is generally regarded as a senolytic (Wang, Lankhorst, and Bernards, 2022), is ineffective in reversing ispinesib resistance by itself. Further, we showed that while saracatinib inhibits EGFR as well as SRC, dasatinib does not; and while dasatinib inhibits phosphorylation of STAT3 at Y705 it does not alter EGFR-mediated phosphorylation at S727. However, combining dasatinib with the EGFR inhibitor erlotinib is as effective as saracatinib in reversing ispinesib resistance. Our results emphasize that whether a drug is a senolytic or not is context dependent, and requires an understanding of how resistance and TIS develop and are maintained on a case-specific basis.

We would predict that alternating dosing of a spindle inhibitor with an appropriate senolytic should also be effective, as our model suggests that the TIS state is relatively stable. While treating with a spindle inhibitor would eliminate a large fraction of proliferating cells, it would still allow TIS cells to block quiescent cells from entering the cell cycle and repopulating the proliferative compartment. Following with a senolytic would then reduce the TIS population, induce quiescent cells to become proliferative, and in the process expand the population of cells that are sensitive to spindle inhibitors. One potential problem with alternating between drugs that inhibit different targets is that this approach may allow the tumor to evolve when the first drug is replaced by the second, due to activation of bypass pathways which neutralize the efficacy of the first drug (Loria *et al*., 2022). However, two features argue against this concern. First, we find that prolonged treatment of GBM cells with ispinesib, alisertib, or volasertib produces a stable phenotype characterized by nuclear atypia and cytoplasmic enlargement. While the large/complex cells from resistant tumors may be able to remain in the cell cycle, they do not appear to revert to a normal mitotic phenotype, which we would have expected if alternative pathways that produce normal mitosis were activated. Second, alternating between ispinesib and saracatinib is no less effective than simultaneously treating with both drugs (**Fig. 7D**). In addition to protecting tumor cells from the cytotoxicity of spindle inhibitors, TIS cells also support tumor progression by suppressing anti-tumor immunity. This point is highlighted by our finding that TIS cells produce activated TGFβ, a well-documented immunosuppressant (Yang *et al*., 2010) (**Fig. 6 I-N**). Thus, although TIS cells may be responsible for slowing tumor growth, they nonetheless enhance tumor lethality in multiple ways. We propose that our approach to understanding how TIS and STAT3 drive resistance may apply to the development of resistance to other cytotoxic/cytostatic therapies, and it should be instructive in designing treatment schemes that both optimize efficacy and reduce toxicity to improve patient outcome in glioblastoma.

## ACKNOWLEDGEMENTS

SSR is supported by NIH grants NS073610, NS118513, NS119714, and CA210910, and by a Translational Adult Glioma Award from the Ben and Catherine Ivy Foundation. PC is supported by NIH grants NS073610 and NS118513. PAS is supported by NIH grant NS118513. RSK is supported by NIH grant NS118513. CTM was supported by K99/R00 Pathway to Independence Award from NIAID (1K99AI175656).

We wish to thank Dr. Justin D. Lathia (Lerner Research Institute of the Cleveland Clinic Foundation) for his gift of L1 cells, Drs. Alfredo Quiñones-Hinojosa and Hugo Guerrero-Cazares (Mayo Clinic Florida) for their gift of GBM612 cells, Dr. Roland Friedel (Mount Sinai School of Medicine) for his gift of SD2 and SD3 cells and for technical advice, and Drs. Loic Deleyrolle (University of Florida) and Terrence Burns (Mayo Clinic Rochester) for helpful discussions.

## AUTHOR CONTRIBUTIONS

*Conception and design:* R. Kenchappa, A. Dovas, N. Zarco, S. Rosenfeld, P. Canoll,

*Development of methodology:* R. Kenchappa, A. Dovas, CT MeyerS. Rosenfeld, P. Canoll

*Acquisition of data:* R. Kenchappa, N. Zarco, A. Dovas, V. de Araujo Farias, A. Haddock, NKH Nagaiah

*Analysis and interpretation of data:* R. Kenchappa, N. Zarco, V. de Araujo Farias, A. Dovas, P.A. Sims, C.T. Meyer, P. Canoll, D. Hambardzumyan, S. Rosenfeld

## STAR METHODS

### Resource Availability

#### Lead contact

Further information and requests for resources and reagents should be directed to and will be fulfilled by the lead contact, Steven S. Rosenfeld.

#### Materials availability

Genetically engineered mouse models and cell lines will be distributed after completion of the relevant Materials Transfer Agreements with the Mayo Clinic.

#### Data and code availability

- Bulk RNA-seq data have been deposited in the Gene Expression Omnibus (GEO) database with sequential accession numbers of GSM8267005-GSM8267016.
- Accession information is available in the Key Resources Table. All other data reported in this paper will be shared by the lead contact upon request.
- This paper does not report any original code.
- Any additional information required to reanalyze the data is available from the lead contact upon request.

### Experimental Model and Subject Detail

#### Mice

All mouse procedures were performed in compliance with the Mayo Clinic Institutional Animal Care and Use Committee guidelines. Homozygous floxed *Trp53* mice (Stock #008462) were obtained from Jackson Laboratory. Studies were performed on equal numbers of male and female mice, ages 8-20 weeks. Animal genotypes were regularly verified via tail snip (TransnetYX, Cordova, TN).

### Method Details

#### Glioma cell line isolation from mouse GBM tumor and culture

The protocol for isolation of tumor cells from *Trp53(−/−)* and *Trp53/Pten(−/−)* murine tumors has been described (Lei *et al*. 2011). Murine mesenchymal (MES) glioblastoma cells (MES1861 and MES4622) which lack expression of Nf1 and Trp53 were maintained in DMEM media with 10% FBS as previously described (Gursel et al., 2011; Reilly et al., 2000), while mouse proneural (PN) glioblastoma cells PN20, PN24 were derived from primary PDGFB-driven glioblastomas generated in Nestin-tva: Cdkn2A knockout mice (Hambardzumyan et al., 2009) and grown in mouse neural stem cell medium (STEMCELL Technologies, 05700 and 05701), supplemented with 20 ng/ml hEGF (Sigma-Aldrich, E9644), 10 ng/ml hFGF (R&D systems, 233-FB-025) and 2 mg/ml heparin (STEMCELL Technologies, 07980). The human L1 primary GBM cell line was cultured and maintained in DMEM+F12 media with 1% N2 supplement (Gibco), 20ng/ml of hEGF (Sigma-Aldrich) and 20ng/ml of hFGF (R&D systems). Human GBM612 primary lines were cultured and maintained in DMEM+F12 media with 1% NeuroPlex supplement (Gemini), 20ng/ml of EGF and 20ng/ml of FGF. Human GBM TS543 line was cultured and maintained in NeuroCult NS-A medium with proliferation supplement (Stem cell technologies), 20ng/ml of EGF, 20ng/ml of FGF and 2ug/ml of heparin. Ispinesib, alisertib or volasertib resistant *Trp53/Pten(−/−)* murine GBM cells were generated by culturing in the presence of 75 nM of ispinesib, 50nM of alisertib or 50nM of volasertib for three weeks, and then maintained at same concentration of these drugs. Ispinesib, alisertib or volasertib resistant human L1, 612 and TS543 GBM cells were generated by culturing in the presence of 25 nM of drug for first week and 50 nM for second and third week, and were maintained in the presence of 50 nM of drug.

#### Dose response curves/cell viability assays

5,000 cells/well were plated in 96-well plates and were allowed to attach for 48 hours. Cells were treated with various doses of ispinesib, alisertib, volasertib, saracatinib, SH5-07 or vehicle for 72 hours and cell viability was measured using CellTiter-Glo (Promega, cat# G9242).

#### Retrovirus production, intracerebral injections and drug treatment

PDGF-IRES-cre retrovirus was generated and injected intracranially according to methods described previously (Lei *et al*., 2011; Kenchappa *et al*., 2020). For the pharmacologic studies mice were treated seven days after retroviral injection with vehicle, alisertib (30 mg/kg by oral gavage, 5 days per week), saracatinib (25 mg/kg by oral gavage, 5 days per week), volasertib (10 mg/kg by intraperitoneal injection for every 4 days), or alisertib + saracatinib (alisertib: 30 mg/kg by oral gavage, 5 days per week; saracatinib: 25 mg/kg by oral gavage, 5 days per week) or volasertib + saracatinib (volasertib: 10 mg/kg by intraperitoneal injection for every 4 days; saracatinib: 25 mg/kg by oral gavage, 5 days per week). For simultaneous vs sequential ispinesib and saracatinib experiments, mice were administered with vehicle, ispinesib (10 mg/kg by intraperitoneal injection every 4 days), ispinesib + saracatinib simultaneously (ispinesib: 10 mg/kg by intraperitoneal injection every 4 days; saracatinib: 25 mg/kg by oral gavage, 5 days per week) or sequentially (alternating cycles of ispinesib 10 mg/kg by intraperitoneal injection for every 4 days for 3 weeks, followed by 1 week of saracatinib 25 mg/kg by oral gavage, 5 days per week). Treatment continued until tumor morbidity.

#### Brain histological analysis

Brains from 4% paraformaldehyde-perfused, GBM-bearing mice were paraffin-embedded as described (Kenchappa et al., 2020). Immunohistochemistry was performed on 5 mm sections using the Discovery ULTRA automated stainer (Ventana Medical Systems). Antigen retrieval was performed using a Tris/borate/EDTA buffer (Discovery CC1), pH 8.0-8.5, for 60 minutes at 95^0^C. Slides were incubated with anti-Ki67 or anti-p21 for 2 hours at room temperature. The antibodies were visualized using biotinylated goat anti-rabbit and rabbit anti-rat secondary and counterstained with hematoxylin and eosin (H&E). Images were captured and analyzed using a ScanScope scanner and ImageScope software (Aperio Technologies).

#### Immunofluorescence microscopy

Drug naïve and resistant GBM cells were grown on glass bottom chamber slides and fixed with ice-cold 4% paraformaldehyde for 20 minutes at room temperature. After washing with PBS three times, cells were permeabilized with 0.1% Triton X-100 PBS, and then washed three more times with PBS. Cells were incubated with 10% donkey serum in PBST (PBS+ 0.1% Tween 20) for 1hr to block non-specific binding of the antibodies, then with Ki67 antibody (MA5-14520, Thermo fisher Scientific) 1:500 in 2% donkey serum in PBST overnight at 4°C. Following washing, a secondary antibody (donkey anti-rabbit IgG-Alexa 488, Molecular Probes, cat #A21206) diluted 1:500 in PBST was applied to cells and incubated 1 hr at room temperature in the dark. Cells were stained for F-actin with Rhodamine Phalloidin (Cytoskeleton Inc, cat #PHDR1, diluted to 100nM final concentration) for 30 minutes. The slides were then mounted with cover slips using Vectashield (#H-1200; Vector Laboratories). Visualization and imaging of the stained cells were conducted using a confocal microscope (Zeiss LSM 880) with a 10x objective lens. The Alexa Fluor 488 settings were employed to detect the Ki67 signal. Concurrently, images were captured to visualize the Rhodamine phalloidin staining utilizing the Rhodamine settings. Images were acquired from three independent experimental replicates to ensure the reliability and reproducibility of the results.

#### Mitochondrial membrane potential assay

To measure the mitochondrial membrane potential in drug naïve and ispinesib resistant mouse GBM cells we used JC-1 (tetraethylbenzimidazolylcarbocyanine iodide; Abcam, cat #ab113850). Drug naïve and ispinesib resistant cells were cultured on chamber slides. After 72 hrs, cells were washed once with 1XPBS and treated with 10µm JC-1 solution in dilution buffer for 15 minutes at 37^0^C, then washed twice with dilution buffer. In some experiments, ispinesib resistant GBM cells were treated with vehicle or 500nM of erlotinib for 48 hr to inhibit EGFR signaling and then added with 10µm of JC-1 solution for 15 minutes. Live cell imaging was conducted to measure green (FITC) and red (TRITC) fluorescence using a Zeiss confocal microscope (LSM 880) equipped with a 10X objective. The imaging environment was meticulously controlled for temperature, CO2 levels, and humidity to ensure optimal conditions for live cell analysis. In most experiments, a single image was acquired from each of four independent experimental replicates. For experiments involving erlotinib treatment, 3-4 images were captured from each of three independent experimental replicates. The total fluorescence from the entire field of view was quantified using ImageJ software.

#### Beta-galactosidase staining of GBM cells

For senescence associated β-galactosidase assay, we used a β-galactosidase staining kit (Cell Signaling Technology, cat#9860). Drug naïve and resistant GBM cells were washed once with 1XPBS and fixed in 1X fixative solution for 15 min at room temperature. Cells were then washed two times with 1XPBS and incubated with β-galactosidase staining solution (pH 6.0) at 37^0^C in a dry incubator without the added CO_2_, and plates were sealed with parafilm to prevent evaporation of the staining medium. After the overnight incubation, cells were washed with 1XPBS and stained images were observed and captured using bright field microscope. In some experiments, ispinesib was withdrawn from ispinesib resistant GBM cells for 24 hours, which were then treated with vehicle or 500nM of saracatinib. 72 hours later cells were washed once with 1XPBS, fixed with 4% paraformaldehyde for 10 minutes, washed with PBS containing 1% BSA, and incubated with fluorescent β-galactosidase staining solution (CellEvent™ Senescence Green Detection Kit, cat# C10850, ThermoFisher Scientific) at 37^0^C in a dry incubator without CO_2_ for 2 hours. Cells were washed with 1XPBS and stained for F-actin and DAPI. Cells were visualized and imaged using confocal microscopy (Zeiss).

#### Mitochondrial respiration assay using the Seahorse XF96 Cell Mito Stress Test

We measured the mitochondrial oxygen consumption rate (OCR) using a Seahorse XFe96 extracellular flux analyzer and a Seahorse XF Cell Mito Stress Test Kit (XF cell mito stress kit, cat#103708-100; XF cell mito stress, cat#103015-100, Agilent Technologies). Drug naïve and ispinesib resistant GBM cells were plated in Seahorse XF Cell Culture Miniplates, treated with vehicle or 500nM of erlotinib for 24 hours and the Cell Mito Stress assay was performed according to the manufacturer’s protocol.

#### Beta-galactosidase vs FSCA and SSCA flow cytometry

Cellular senescence and cellular size and complexity were evaluated in *Trp53/Pten*(−/−) murine GBM and L1 human GBM cell lines. Senescence was assessed through flow cytometry employing the CellEvent™ Senescence Green Flow Cytometry Assay Kit. Drug-resistant cells were generated by exposing *Trp53/Pten*(−/−) cells to 75 nM of ispinesib, 50 nM of alisertib, or 50 nM volasertib over a three-week period, followed by maintenance at the same concentrations. Resistant L1 human GBM cells were induced by culturing them in the presence of 25 nM of the respective inhibitors during the first week and 50 nM during the subsequent weeks, with maintenance at 50 nM. For flow cytometric analysis, 500,000 cells were harvested, fixed with 2% PFA, and incubated with a 1:500 dilution of the CellEvent™ Senescence Green probe for 2 hours at 37°C in the dark under an environment devoid of CO_2_. Data acquisition was carried out using a Cytoflex flow cytometer, measuring β-gal-positive cells. Drug-naïve and non-stained cells were utilized as controls to establish forward and side scatter thresholds and to identify negative and positive β-gal cell populations. Data analysis was performed using FlowJo v10.9 software. All experiments were conducted in triplicate.

#### Beta-galactosidase vs Ki67 flow cytometry

To evaluate cell senescence and proliferation, a flow cytometry-based assay was conducted utilizing *Trp53/Pten*(−/−) murine GBM cells under various experimental conditions. The cells were stratified into the following groups: drug-naïve, Ispinesib-resistant, and Ispinesib-resistant treated for 2 days with 500 nM saracatinib. Fluorescent staining of β-gal activity was performed according to the protocol described abover. Following β-gal staining, the cells underwent immunofluorescence staining to assess cell proliferation via Ki67 expression using a 1:200 dilution of the primary antibody. A lexa Fluor 594-labeled secondary antibody was utilized to visualize Ki67 positive cells. Sample data were collected using a Cytoflex flow cytometer to quantify β-gal and Ki67 positive cells. Drug-naïve and non-stained cells were utilized as control to set up the flow cytometry analysis and differentiate negative and positive populations for β-gal and Ki67. Data analysis was performed using FlowJo software version 10.9 to assess the proportions of senescent and proliferative cells within each experimental group. The study was replicated across three independent experiments.

#### H2B-GFP lentivirus construction

To generate an inducible H2B-GFP expressing lentiviral construct, a molecular cloning process was employed. Using SnapGene software, the lentiviral vector was designed by incorporating the H2B and GFP (pLenti Lifeact-EGFP BlastR, Addgene #84383) sequences into the plasmid pCW-Cas9 (Addgene #50661). cDNA encoding the histone H2B protein was amplified from SD2 cells (a generous gift of Dr. Roland Friedel, Mount Sinai School of Medicine) following induction with doxycycline. All the PCR reactions were carried out using high-fidelity DNA polymerase (Takara), and specific primers that are detailed in the Key Resources table. The amplified PCR products were purified via agarose gel extraction. The ligation was conducted using NEBligase, followed by transformation into NEB Stable competent cells. Selection of transformants was performed on ampicillin-containing agar plates, and ampicillin-resistant colonies were expanded for plasmid extraction via miniprep. The integrity of the assembled lentiviral construct was confirmed through enzymatic restriction digestion. Functional validation of the construct was achieved by transfecting HEK cells and inducing expression with doxycycline to ascertain the inducible expression of the H2B-GFP protein.

#### H2B-GFP lentivirus production

To generate H2B-GFP lentivirus, HEK-293T cells were cultured under optimal conditions until they reached 80% of confluency for transfection. The cells were subsequently co-transfected with a recombinant lentiviral vector encoding the H2B-GFP protein, along with the psPAX2 and pMD2.G packaging plasmids utilizing Lipofectamine 3000, according to the manufacturer’s protocol, in Opti-MEM reduced serum medium. After six hours of transfection the medium was replaced with complete growth medium to support cell viability and promote viral production. Viral supernatants were harvested at two sequential time points: 24- and 48-hours post-transfection. These supernatants were combined to maximize viral yield. Following collection, the lentiviral particles were concentrated using the Lenti-X concentrator system as per the manufacturer’s guidelines. The concentration of viral particles was quantified using a p24 ELISA assay.

#### H2B-GFP transduction and selection of L1 and Trp53/Pten(−/−) cells

To investigate the proliferative behavior of GBM cells, we employed two distinct cell lines: L1 (*human GBM*) and *Trp53/Pten*(−/−) (*murine GBM*). These lines were genetically engineered to express H2B-GFP. The transduction process was carried out using a lentiviral vector encoding the H2B-GFP construct, described above. Cells were incubated with the viral particles in complete medium containing 4 μg/mL polybrene at an MOI of 50 for 72 hours. After transduction, cells were selected for stable integration of the H2B-GFP construct with 1.5 μg/mL puromycin for two weeks and selection by FACS with the Aria cytometer. The expression and functionality of H2B-GFP were validated through fluorescence microscopy after the induction of with 2 μg/mL doxycycline in complete medium for 48 hours.

#### H2B-GFP fluorescence decay studies

H2B-GFP fluorescence decay was assessed using lentiviral transduced L1 human GBM cells and *Trp53/Pten*(−/−) murine GBM cells, both expressing inducible H2B-GFP protein. To generate resistant cell lines, cells were subjected to ispinesib, alisertib, or volasertib treatment for three weeks, as described above. Induction of H2B-GFP expression was achieved by adding 2 μg/mL doxycycline to the culture medium for 48 hours. Following induction, the medium was replaced with fresh medium, and the decay of GFP fluorescence intensity was monitored over a 24-day period at various time points using a Zeiss LSM 880 confocal microscope. Quantification of fluorescence in the confocal images was conducted using ImageJ software. Data analysis was performed with GraphPad software.

#### Western blots

Cells were scraped and incubated in lysis buffer (50 mM Tris HCl at pH 7.40, 150 mM NaCl, 1 mM EDTA, 1.0% Nonidet P-40, and a mixture of protease and phosphatase inhibitors), on ice from 30 minutes. Debris was removed by centrifugation for 10 minutes at high speed at 4°C, and cleared lysates were run on SDS/PAGE and transferred to polyvinylidene difluoride membranes. Membranes were blocked in 5% non-fat dry milk in TBS + 0.1% Tween 20 for 1 hour at room temperature, incubated with primary antibody in blocking solution for overnight at 4°C, followed by secondary antibody for 1 hour at room temperature, and developed using an enhanced chemiluminescence solution.

#### Drug synergy

Synergy was calculated using the MuSyC algorithm as previously described (Wooten et al., 2021; Meyer et al., 2019). MuSyC quantifies two types of drug synergy, synergistic potency and synergistic efficacy, both relating to geometric transformations of the dose response surface, which are analogous to the transformations in the 1D Hill equation for potency (horizontal shift in the EC_50_) and efficacy (vertical shift in E_max_). Synergy was calculated by fitting a dose-response surface relating the drug’s effect to the concentrations of drug 1 and drug 2. To help the growth-rate-based fits converge, the bounds for E_0_ (effect with no drug) and E_1_, E_2_, E_3_ (maximal effects of drug 1, 2, and the combination) were set to [0.005, 0.002] and [0.01, −0.01], respectively.

#### STAT3 shRNA encoding lentiviral production and transduction

Knockdown of STAT3 in murine mesenchymal GBM cells was achieved *via* lentiviral infection with shRNA-encoding constructs. The lentiviral plasmid vector pLKO.1-puro based shRNA clones and control shRNA vector were purchased from Sigma-Aldrich (St Louis, MO, USA). The following constructs were used for murine GBM cells in these studies: Non Targeting control (SHC002); STAT3 sh-RNA [TRCN0000071454 (STAT3-shRNA-1), TRCN0000071456 (STAT3-shRNA-2). Each of the pLK0.1 targeting constructs were co-transfected with psPAX2 and pMD plasmids into HEK-293T cells via Lipofectamine 3000 transfection agent (Life Technologies, catalog # 11668027) in serum-free medium. After 8 hours of transfection, the viral particle-containing medium was removed and replaced with fresh complete medium. Transfected cells were then grown in DMEM media containing 10% FBS for 48 hours at 37°C, 5% CO2. Media containing virus was harvested and centrifuged for 10 mins in a clinical specimen centrifuge and then filtered through a 0.45 μm filter. Lentiviral particles were concentrated using LentiX-Concentrator reagent (Takara Bio USA) and the viral titer was determined using a p24 ELISA kit (Clontech). Mouse glioma cells were infected by incubating with virus containing media (at 10 MOI of virus and 4μg/mL of polybrene (Sigma-Aldrich)) overnight. Cells were selected for positive shRNA infection using puromycin (0.5ug/ml) for seven days and maintained in 0.1ug/mL puromycin containing media, and then effect of STAT3 knockdown on cell viability was measured.

#### Bulk RNA-seq data acquisition and analysis

RNA sequencing was performed at the Columbia Sulzberger Genome Center. Total RNA from three independent biological replicates (naïve and ispinesib resistant cells) was isolated using the RNAqueous phenol-free total RNA isolation kit (Ambion, Life Technologies, Grand Island, NY) and DNA contamination in isolated RNA was removed by DNase treatment using TURBO DNA-free™ kit (Ambion, Life Technologies, CA). All samples had an RNA Integrity Number greater than 7.6, as assessed using Agilent Bioanalyzer. Libraries were prepared using the Illumina TruSeq RNA Library Prep Kit v2 and 20 million paired-end, 75 bp reads were acquired on an Element Aviti sequencer at Columbia Genome Center. Reads were pseudoaligned to a kallisto mouse transcriptome index (GRCm38.p6) using kallisto (0.44.0), and differential gene expression analysis was determined using DESeq2. Gene Set Enrichment Analysis (GSEA) was performed on the desktop version of GSEA (v4.1.0), using Hallmarks (Liberzon *et al*., 2015) and the Verhaak_Glioblastoma._mesenchymal gene sets from the Molecular Signatures Database (MSigDB).

### Quantification and Statistical Analysis

#### Statistical analysis

For *in vitro* studies, a two tailed t test or one-way ANOVA was used to calculate p values with statistical significance at p<0.05. For survival studies, statistical significance was determined using a log rank test, and significance was set at p<0.05.

## KEY RESOURCE TABLE

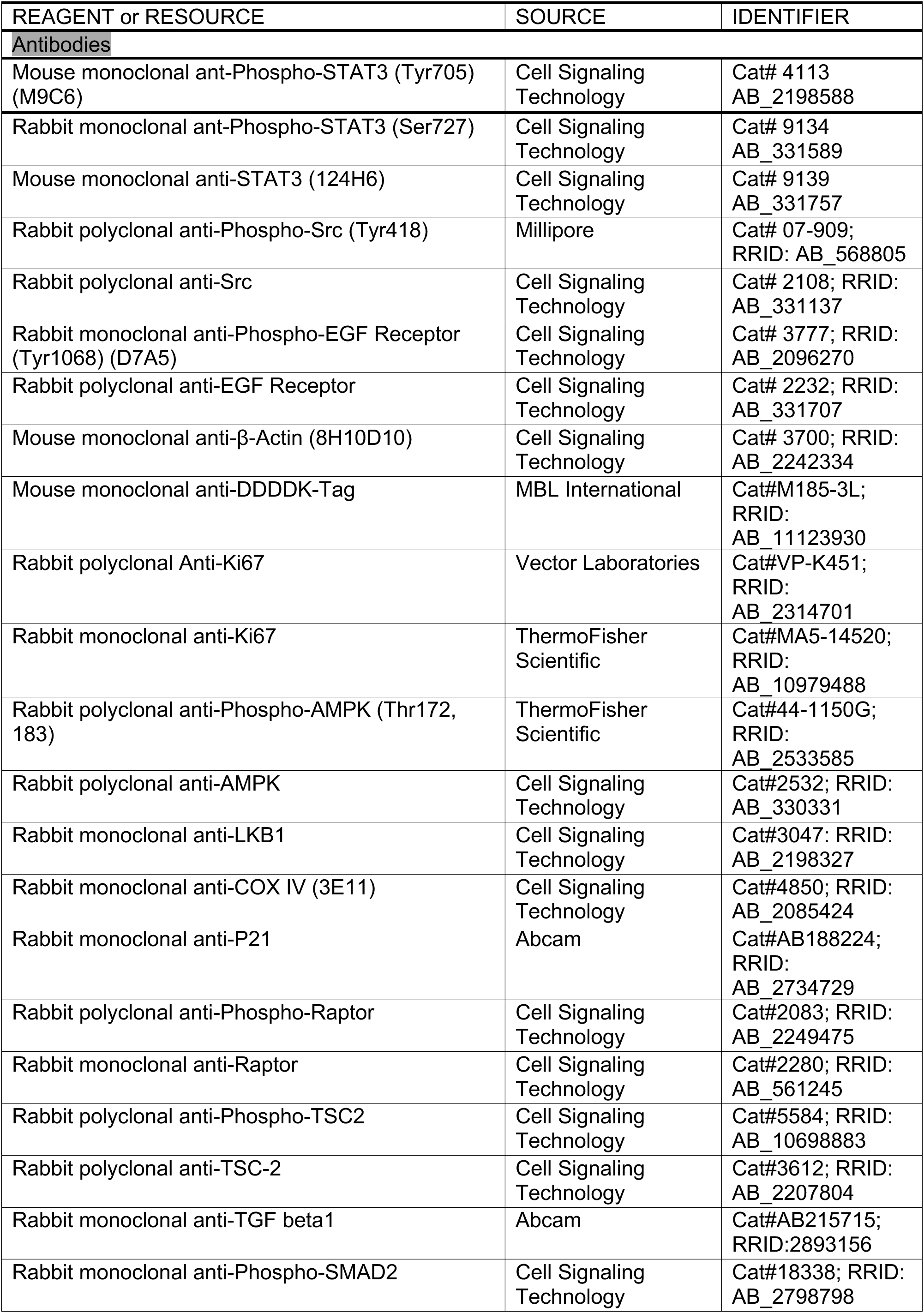
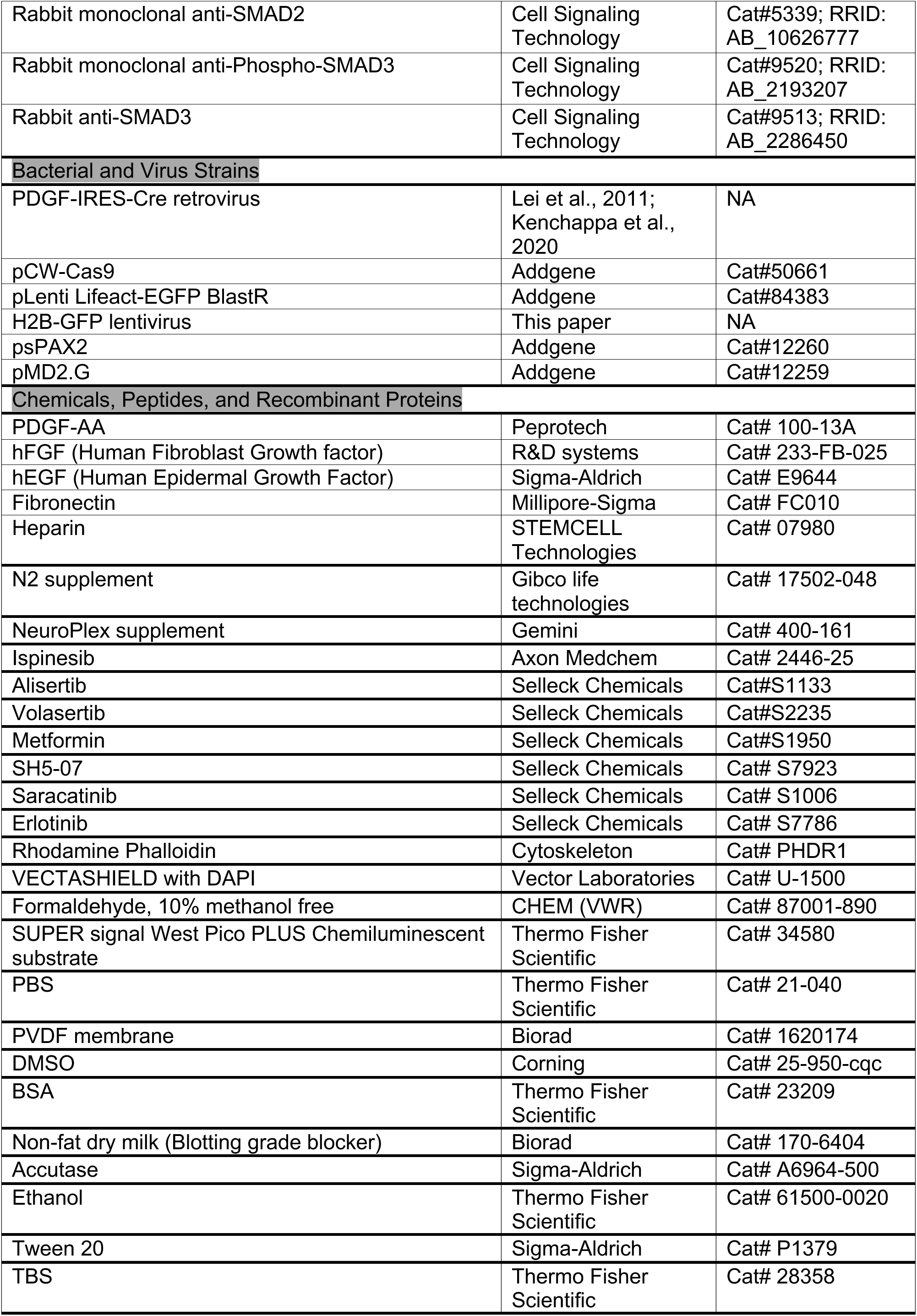
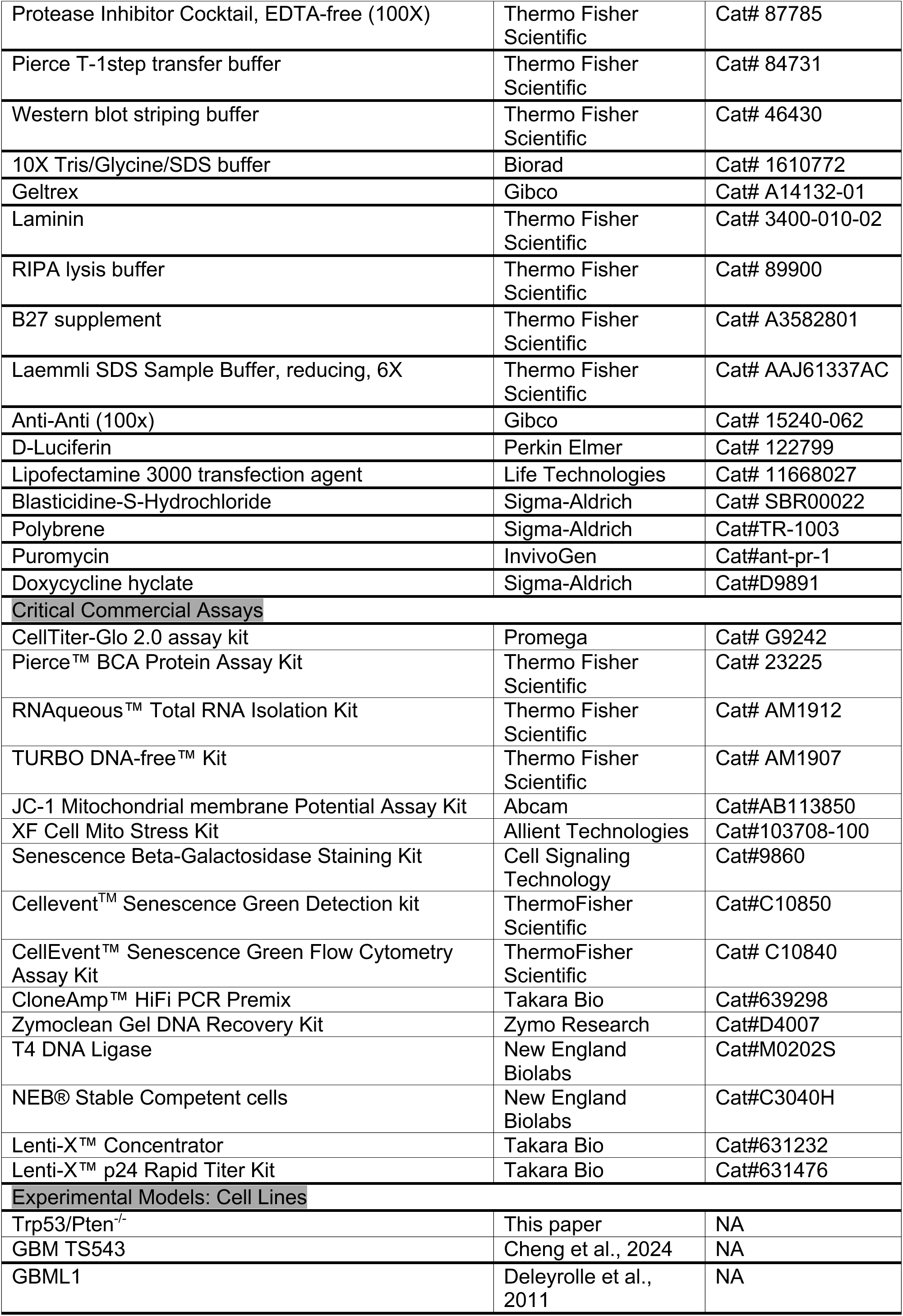
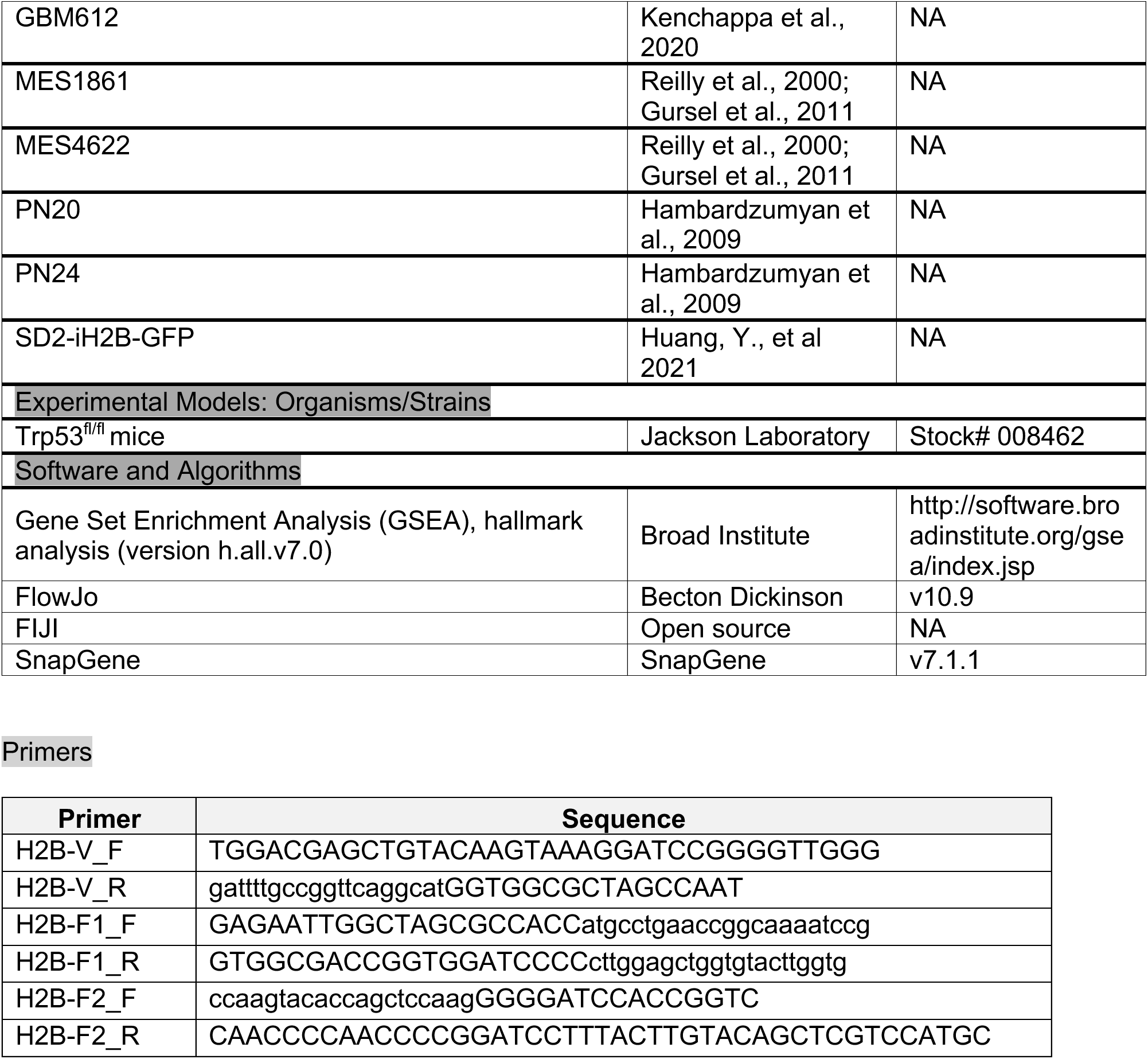

## SUPPLEMENTARY TABLES AND FIGURES

**Table S1:**
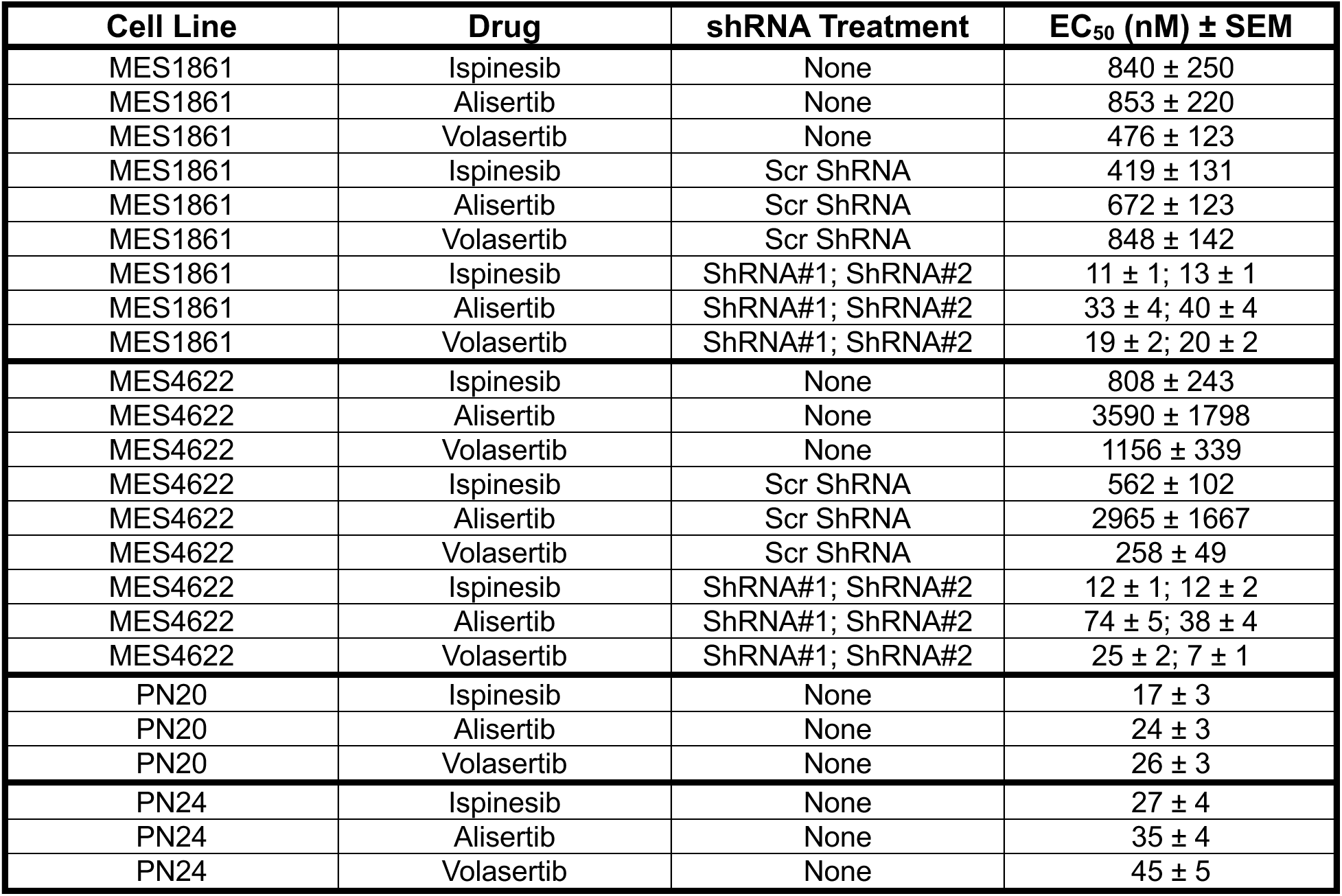
Anti-Mitotic EC_50_ Values for MES1861, MES4622, PN20, and PN24 Cell Lines.

**Table S2:**
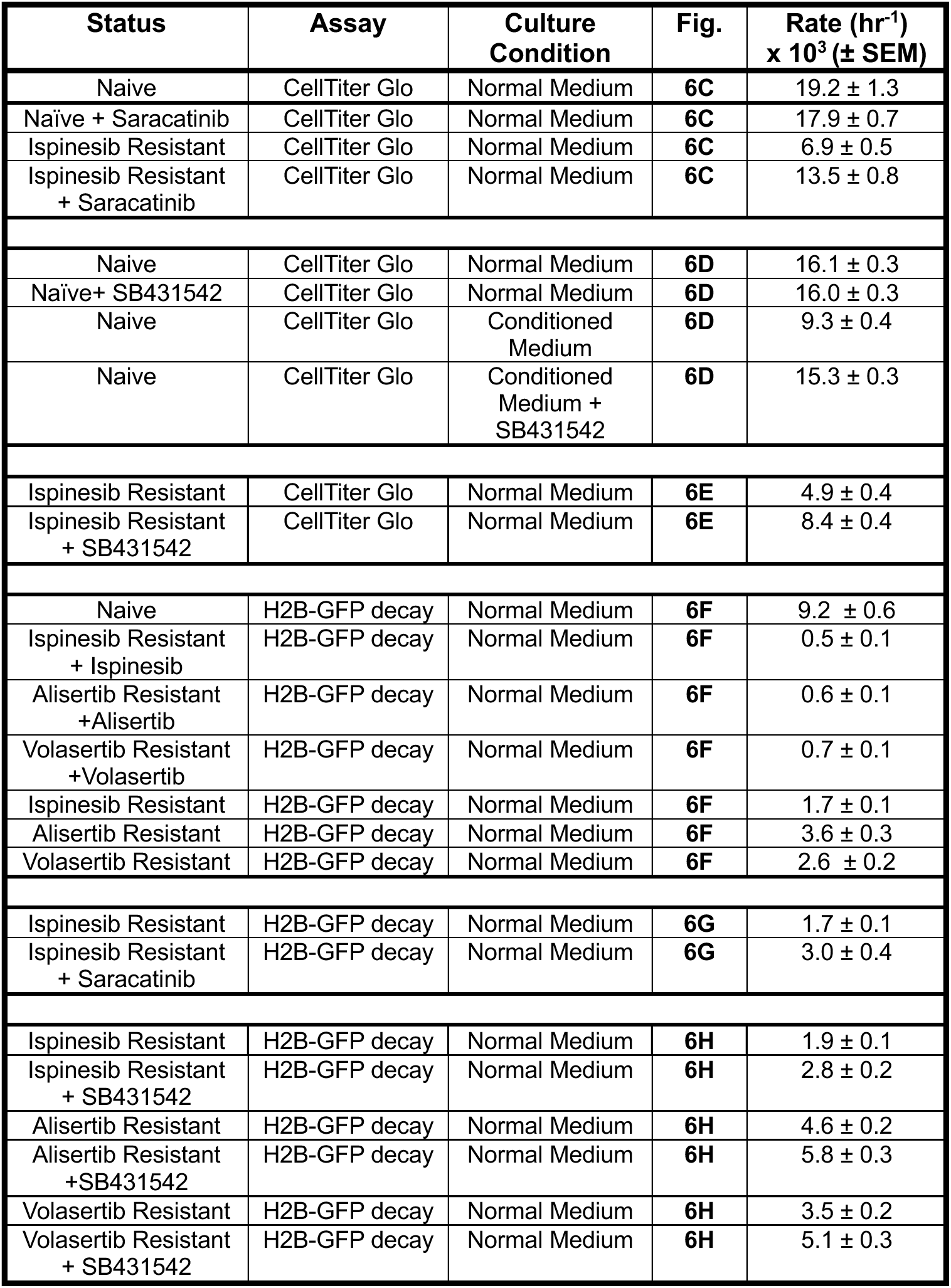
Rate Constants for Proliferation of *Trp53/Pten*(−/−) Cells.

**Table S3:**
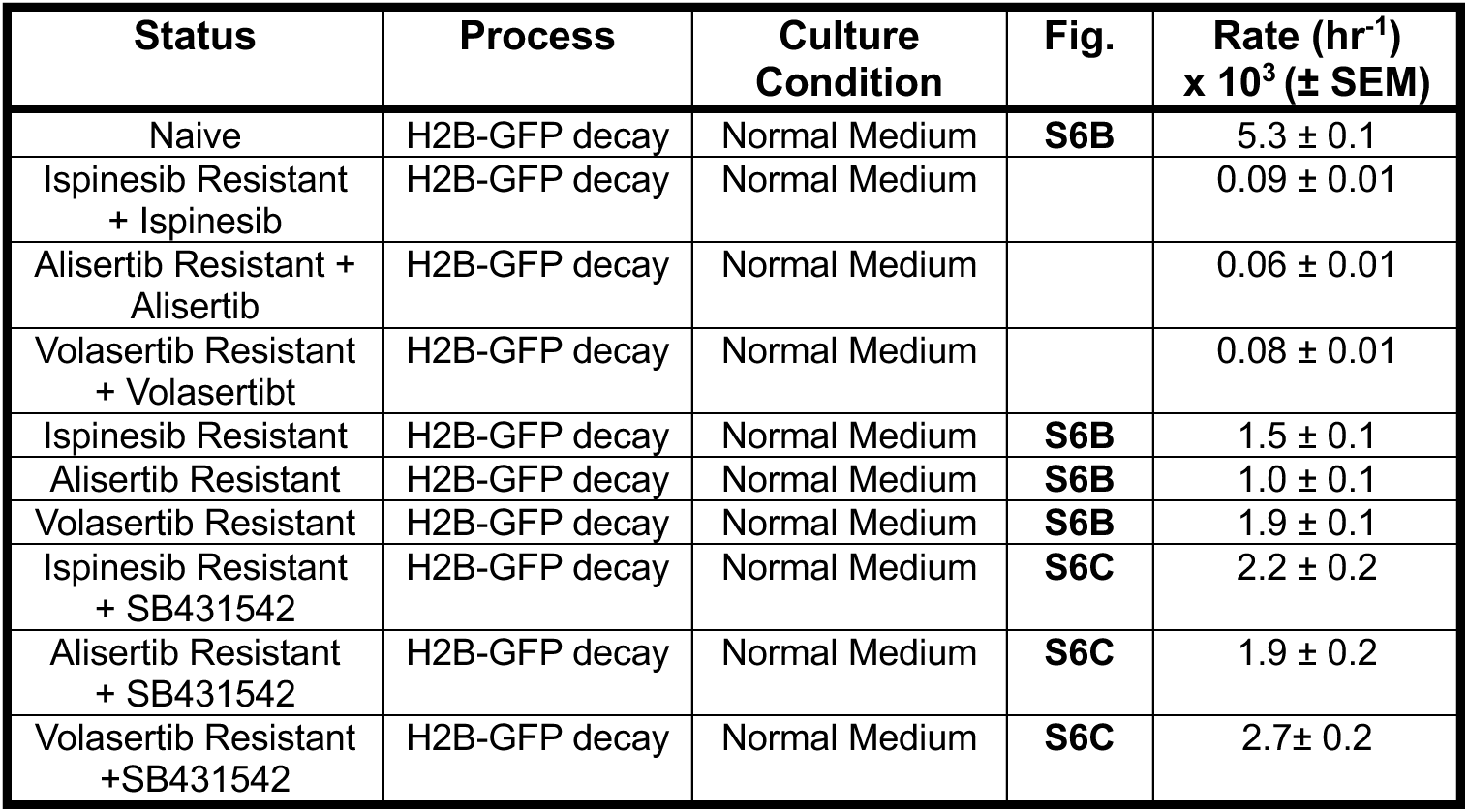
Rate Constants for L1 Cells.

**Fig. S1:**
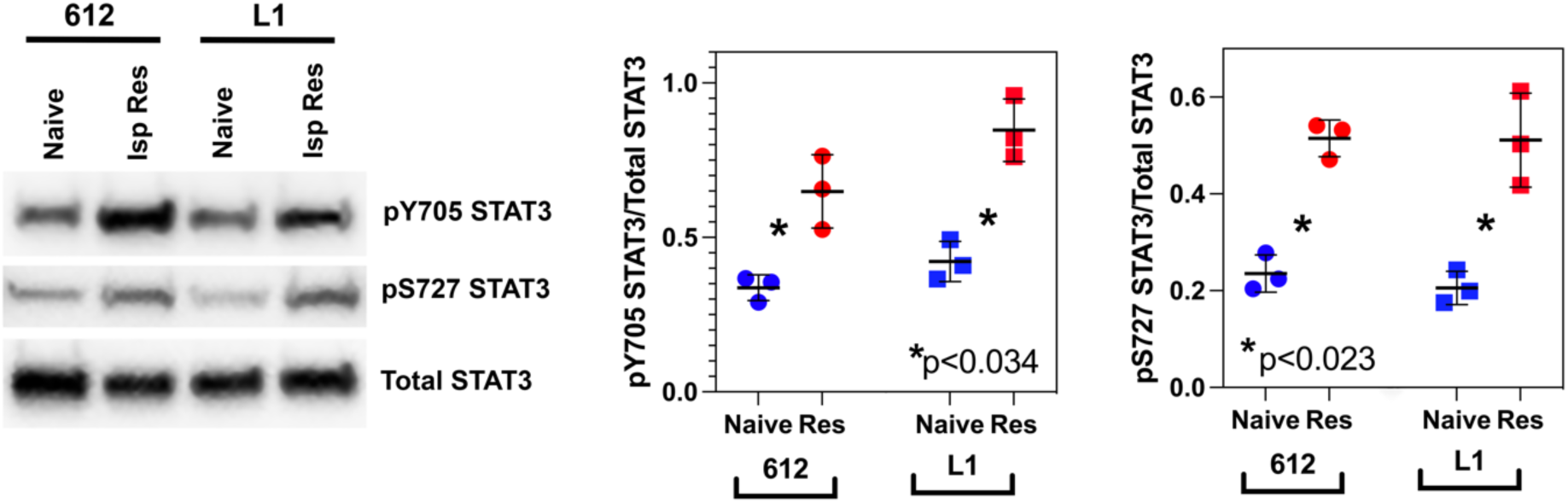
Western blots for pY705 STAT3, pS727 STAT3, and total STAT3 in human 612 and L1 GBM cell lines. For both lines, ispinesib resistance is associated with a statistically significant increase in both phosphorylated species, compared to drug naïve cells.

**Fig. S2:**
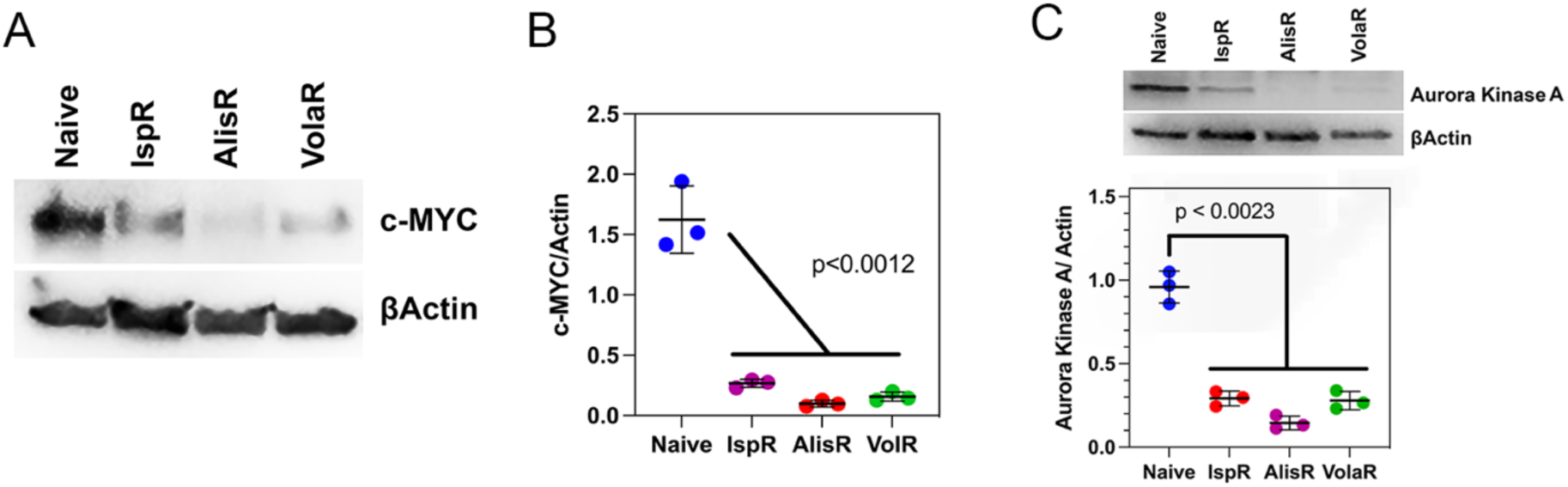
Resistance to spindle inhibitors downregulates c-MYC by 8-10-fold (**A,B**) and downregulates expression of Aurora Kinase A 3-4 fold (**C**).

**Figure S3:**
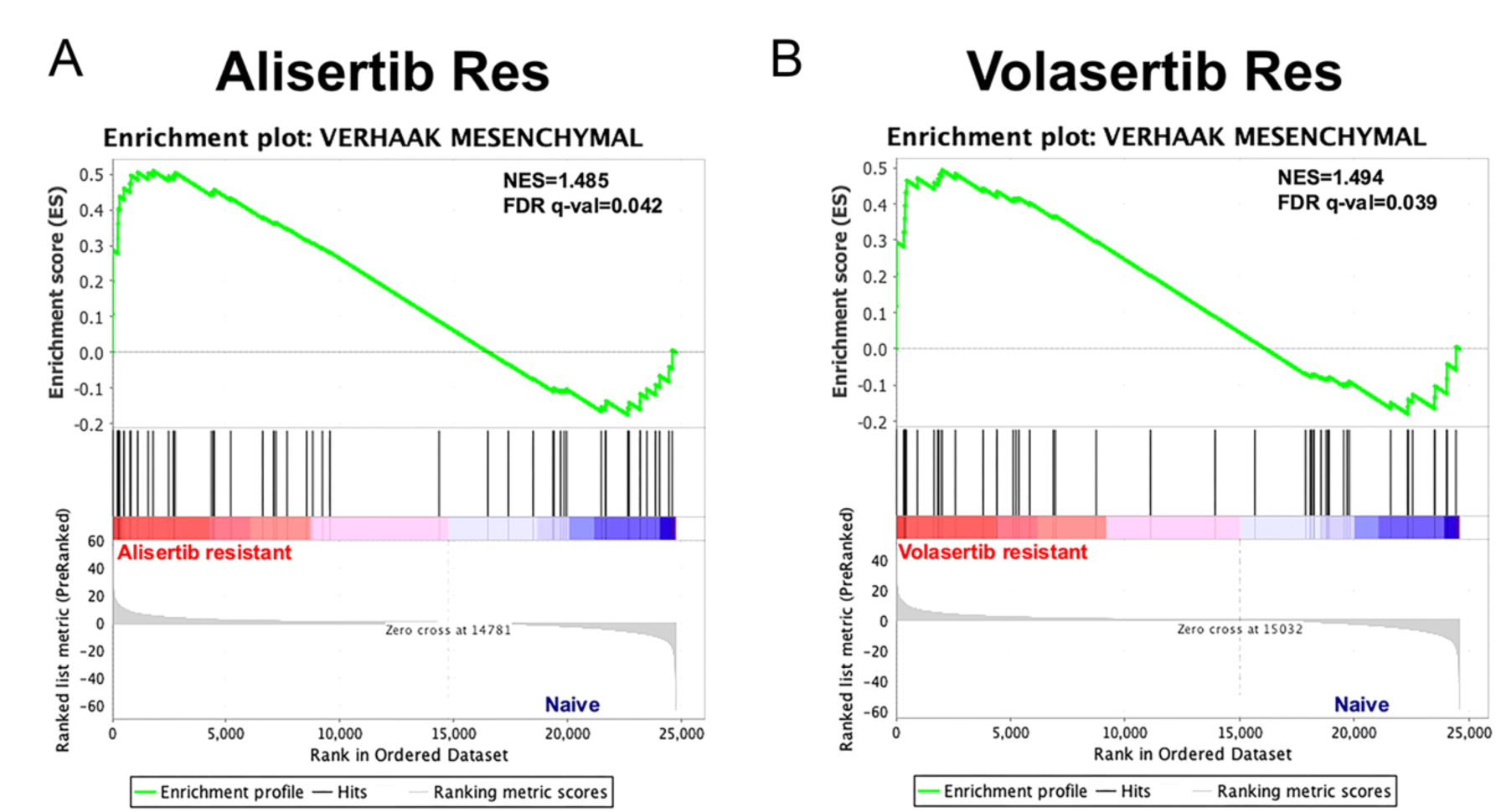
Gene set enrichment analysis of alisertib (A) and volasertib (B) resistant *Trp53/Pten*(−/−) GBM cells. Although the *Trp53/Pten*(−/−) murine GBM line has a strong proneural transcriptional signature, resistance to these two spindle inhibitors, as in the case of ispinesib (Kenchappa *et al*., 2022) produces a proneural→mesenchymal transcriptional shift.

**Figure S4:**
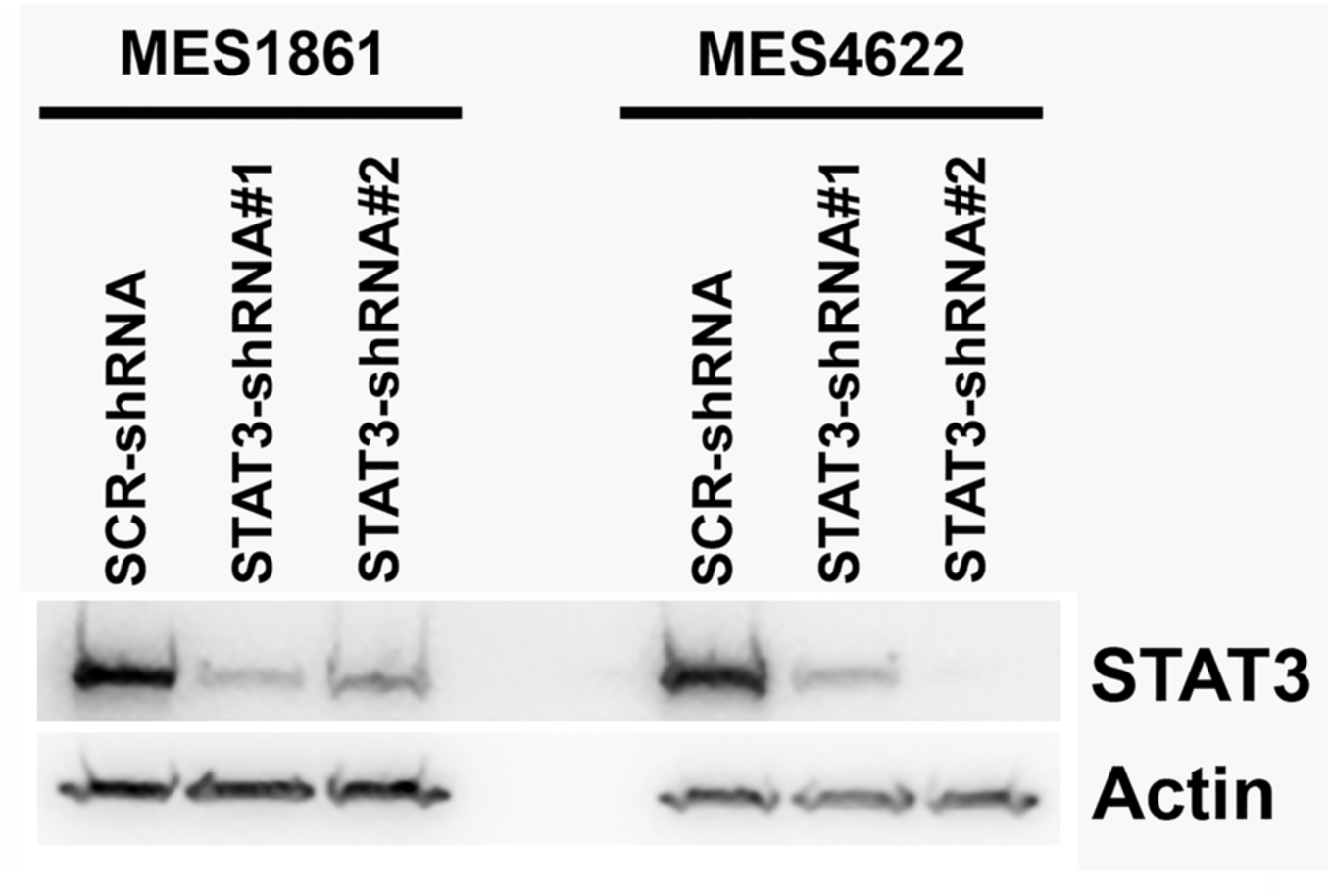
Western blots of STAT3 confirm that both targeting shRNAs reduce STAT3 levels by >90%.

**Figure S5:**
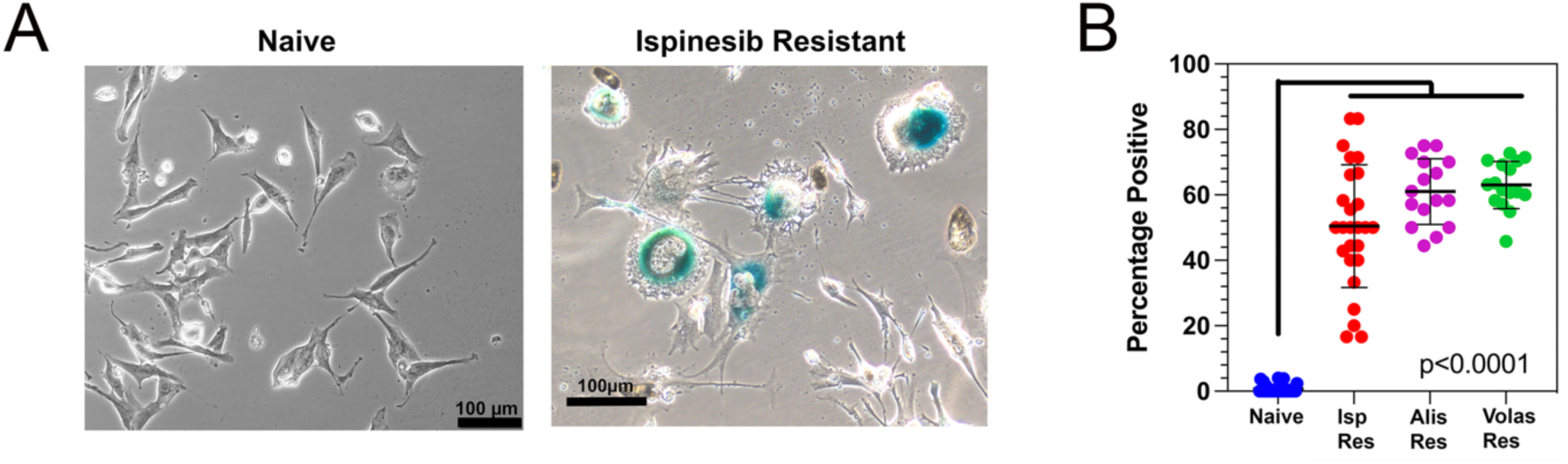
(**A**). X-Gal staining of drug naïve (*left*) and ispinesib resistant (*right*) *Trp53/Pten*(−/−) cells. (**B**). Quantitation of β-gal positivity demonstrates 55-65% positivity in ispinesib (*red*), alisertib (*magenta*), and volasertib (*green*) resistant cells.

**Figure S6:**
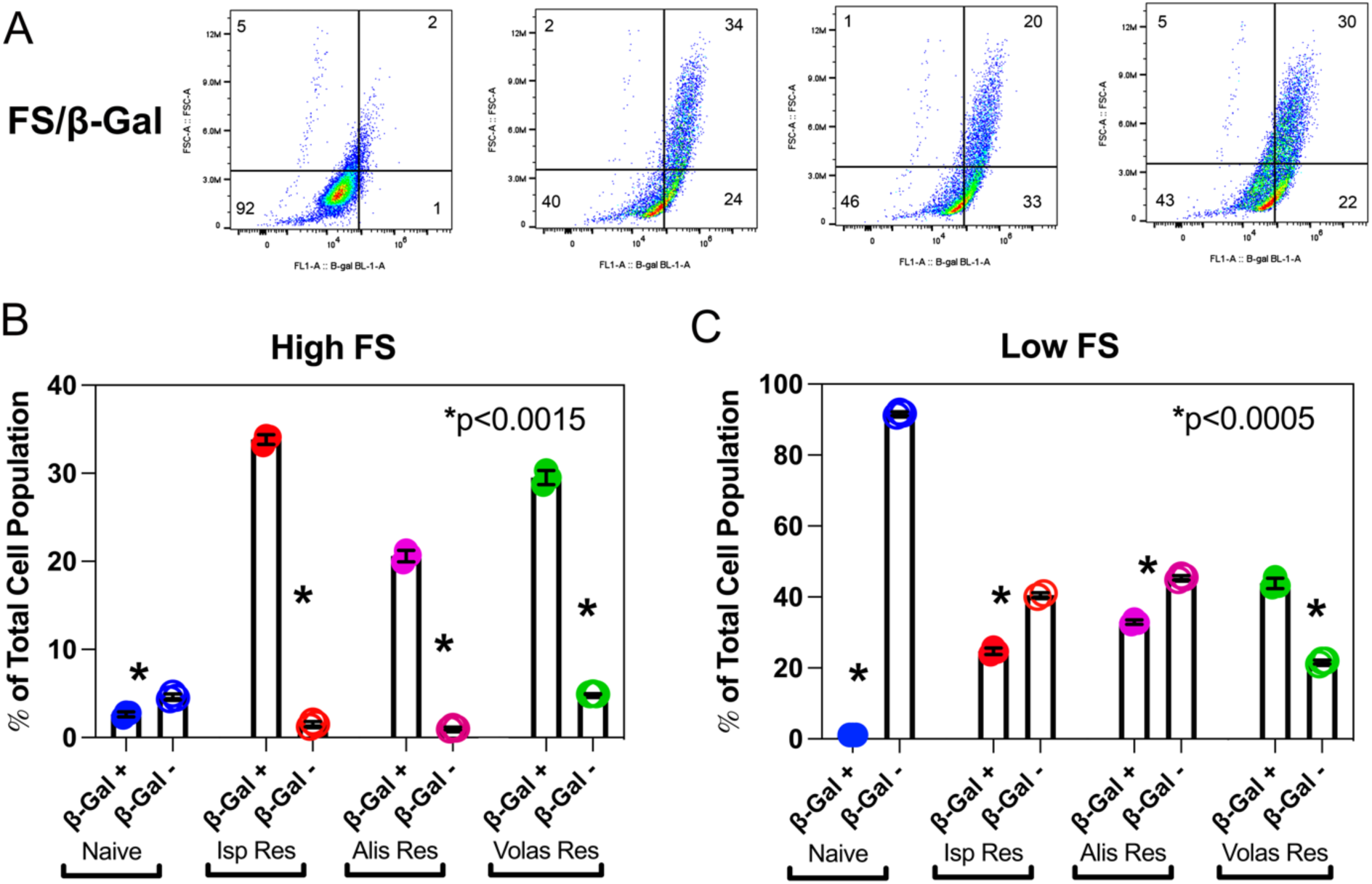
(**A**). Naïve and spindle inhibitor resistant *Trp53/Pten*(−/−) cells were stained with the fluorescent β-gal substrate FDGlu and subjected to flow cytometry to measure forward scatter *versus* fluorescence. A substantial fraction of the resistant cells demonstrate both high forward scatter and β-gal activity. Numbers in each quadrant represent the mean percentages of the total signal. (**B**). Bar plot depicting for naïve and resistant cells the fraction of the total cell population that demonstrates both high forward scatter and either positive or negative FDGlu fluorescence. (**C**). Bar plot depicting for naïve and resistant cells the fraction of the total cell population that demonstrates both low forward scatter and either positive or negative FDGlu fluorescence.

**Figure S7:**
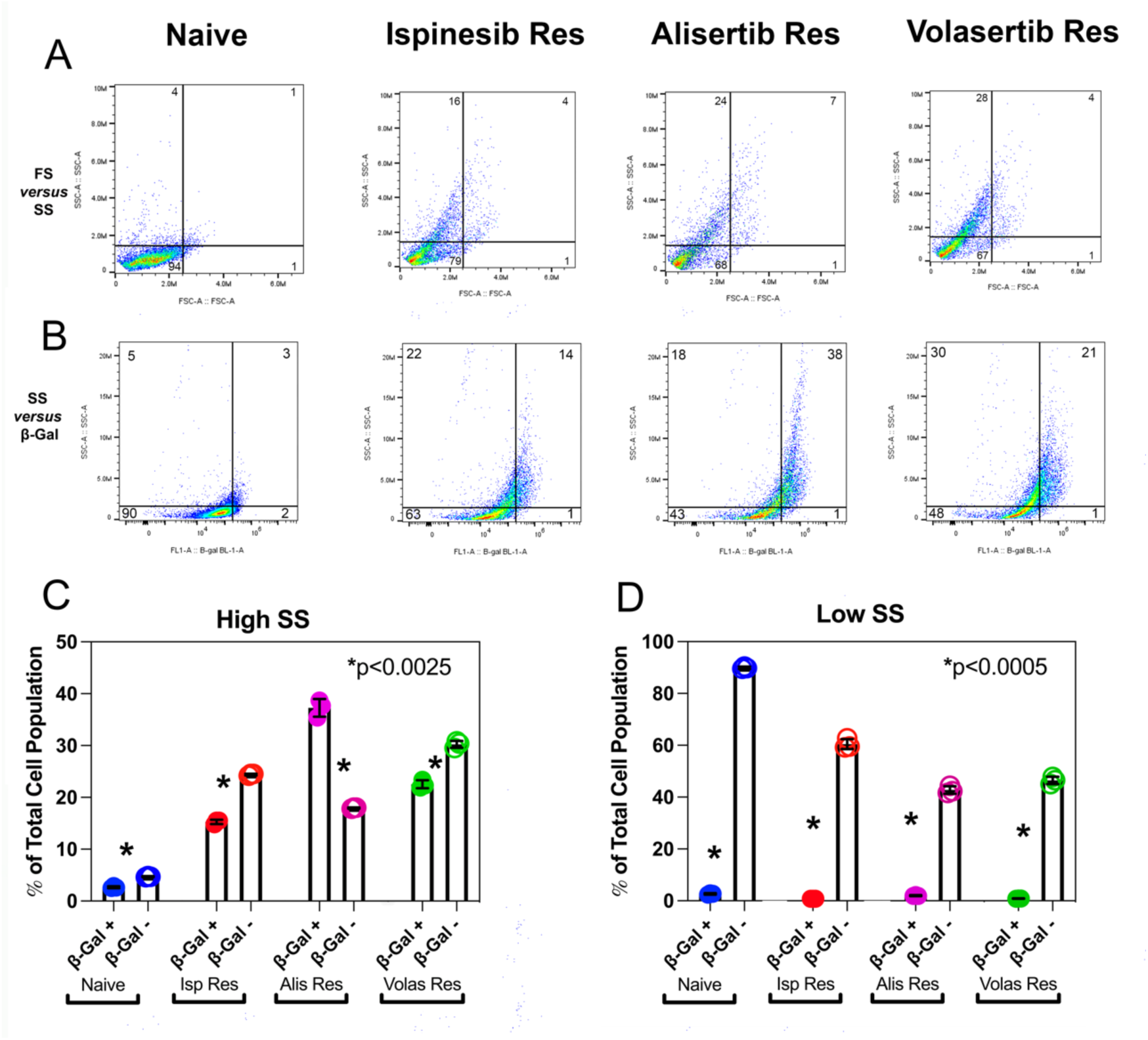
(**A**). Forward versus side scatter flow cytometry of L1 human GBM cells shows that while drug naïve cells cluster in a region of low forward and side scatter, resistance to each of the three spindle inhibitors generates a broad distribution of cells that include a 4-5 fold increase in cells with a high side scatter profile. Numbers in each quadrant represent the mean percentages of the total signal. (**B**). Naïve and resistant cells were stained with the fluorescent β-gal substrate FDGlu and subjected to flow cytometry to measure side scatter *versus* fluorescence. A substantial fraction of the resistant cells demonstrate both high side scatter and β-gal activity. Numbers in each quadrant represent the mean percentages of the total signal. (**C**). Bar plot depicting for naïve and resistant cells the fraction of the total cell population that demonstrates both high side scatter and either positive or negative FDGlu fluorescence. (**D**). Bar plot depicting for naïve and resistant cells the fraction of the total cell population that demonstrates both low side scatter and either positive or negative FDGlu fluorescence.

**Figure S8:**
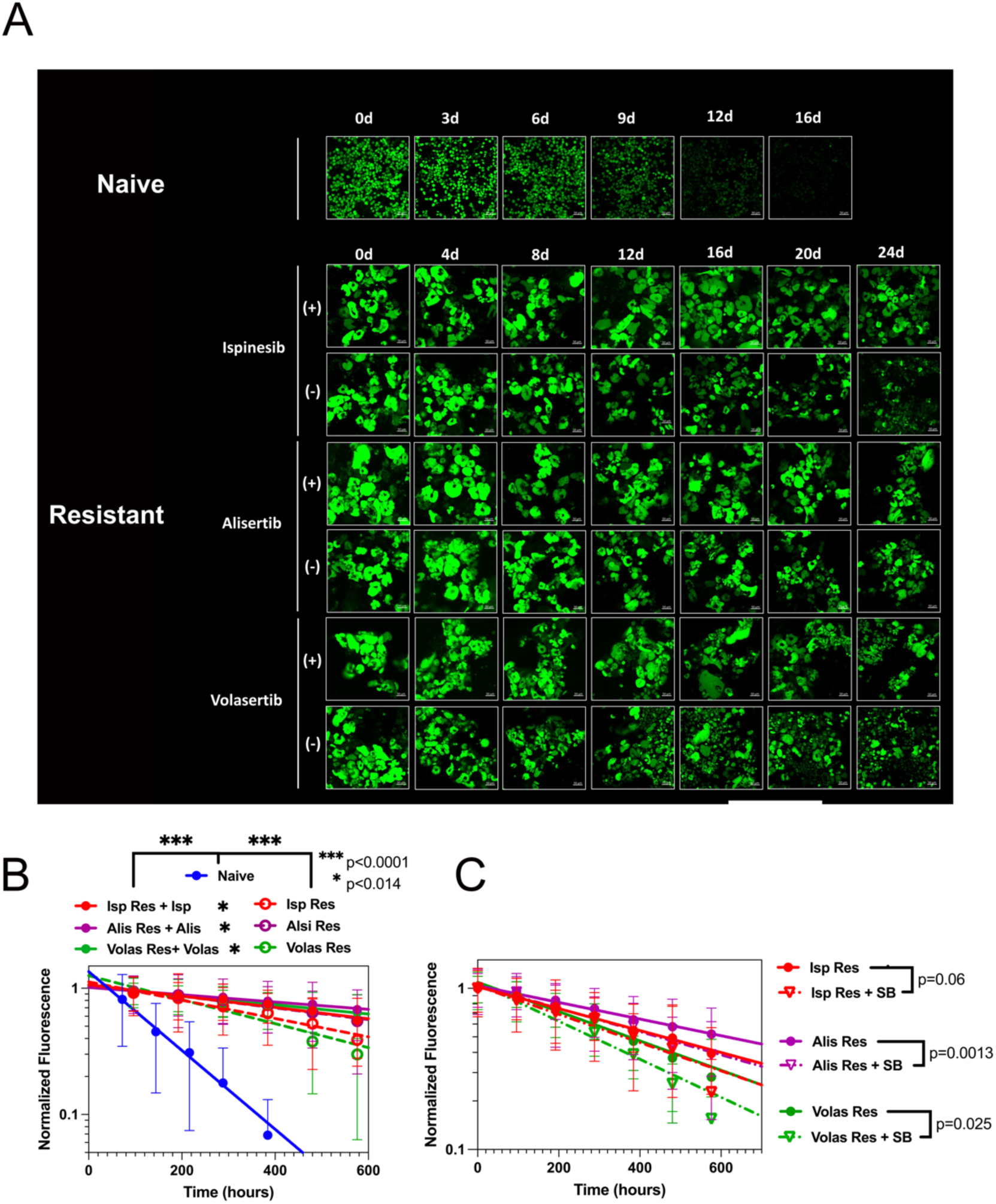
(**A**). Fluorescence micrographs of naïve and spindle inhibitor resistant human L1 GBM cells transfected with doxycycline inducible H2B-GFP. Cells were pulsed with doxycycline, and fluorescence intensity was measured after doxycycline removal, both in the presence (+) and absence (-) of spindle inhibitor. (**B**). Semi logarithmic plot of fluorescence decay of drug naïve (*blue*), ispinesib (red), alisertib (*magenta*), and volasertib (*green*) resistant L1 cells in the presence (*solid circles*) and absence (*open circles*) of spindle inhibitor. Corresponding rate constants from single exponential decays (*lines*) are summarized in **Table S3**. Proliferation of ispinesib (*red*), alisertib (*magenta*), and volasertib (*green*) resistant L1 cells in the absence of spindle inhibitor is accelerated 45-90% by addition of the TGFβ receptor inhibitor SB431542 (**Table S3**).

