## Supplemental Tables and Figures for "Resistance to Spindle Inhibitors in Glioblastoma Depends on STAT3 and Therapy Induced Senescence"

### SUPPLEMENTARY TABLES AND FIGURES

#### Table S1: Anti-Mitotic $EC_{50}$ Values for MES1861, MES4622, PN20, and PN24 Cell Lines

| Cell Line | Drug | shRNA Treatment | $EC_{50}$ (nM) ± SEM |
|---|---|---|---|
| MES1861 | Ispinesib | None | 840 ± 250 |
| MES1861 | Alisertib | None | 853 ± 220 |
| MES1861 | Volasertib | None | 476 ± 123 |
| MES1861 | Ispinesib | Scr ShRNA | 419 ± 131 |
| MES1861 | Alisertib | Scr ShRNA | 672 ± 123 |
| MES1861 | Volasertib | Scr ShRNA | 848 ± 142 |
| MES1861 | Ispinesib | ShRNA#1; ShRNA#2 | 11 ± 1; 13 ± 1 |
| MES1861 | Alisertib | ShRNA#1; ShRNA#2 | 33 ± 4; 40 ± 4 |
| MES1861 | Volasertib | ShRNA#1; ShRNA#2 | 19 ± 2; 20 ± 2 |
| MES4622 | Ispinesib | None | 808 ± 243 |
| MES4622 | Alisertib | None | 3590 ± 1798 |
| MES4622 | Volasertib | None | 1156 ± 339 |
| MES4622 | Ispinesib | Scr ShRNA | 562 ± 102 |
| MES4622 | Alisertib | Scr ShRNA | 2965 ± 1667 |
| MES4622 | Volasertib | Scr ShRNA | 258 ± 49 |
| MES4622 | Ispinesib | ShRNA#1; ShRNA#2 | 12 ± 1; 12 ± 2 |
| MES4622 | Alisertib | ShRNA#1; ShRNA#2 | 74 ± 5; 38 ± 4 |
| MES4622 | Volasertib | ShRNA#1; ShRNA#2 | 25 ± 2; 7 ± 1 |
| PN20 | Ispinesib | None | 17 ± 3 |
| PN20 | Alisertib | None | 24 ± 3 |
| PN20 | Volasertib | None | 26 ± 3 |
| PN24 | Ispinesib | None | 27 ± 4 |
| PN24 | Alisertib | None | 35 ± 4 |
| PN24 | Volasertib | None | 45 ± 5 |

**Table S2: Rate Constants for Proliferation of *Trp53/Pten*(-/-) Cells**

| Status | Assay | Culture Condition | Fig. | Rate ($hr^{-1}$) x $10^3$ (± SEM) |
|---|---|---|---|---|
| Naive | CellTiter Glo | Normal Medium | **6C** | 19.2 ± 1.3 |
| Naïve + Saracatinib | CellTiter Glo | Normal Medium | **6C** | 17.9 ± 0.7 |
| Ispinesib Resistant | CellTiter Glo | Normal Medium | **6C** | 6.9 ± 0.5 |
| Ispinesib Resistant + Saracatinib | CellTiter Glo | Normal Medium | **6C** | 13.5 ± 0.8 |
| Naive | CellTiter Glo | Normal Medium | **6D** | 16.1 ± 0.3 |
| Naïve+ SB431542 | CellTiter Glo | Normal Medium | **6D** | 16.0 ± 0.3 |
| Naive | CellTiter Glo | Conditioned Medium | **6D** | 9.3 ± 0.4 |
| Naive | CellTiter Glo | Conditioned Medium + SB431542 | **6D** | 15.3 ± 0.3 |
| Ispinesib Resistant | CellTiter Glo | Normal Medium | **6E** | 4.9 ± 0.4 |
| Ispinesib Resistant + SB431542 | CellTiter Glo | Normal Medium | **6E** | 8.4 ± 0.4 |
| Naive | H2B-GFP decay | Normal Medium | **6F** | 9.2 ± 0.6 |
| Ispinesib Resistant + Ispinesib | H2B-GFP decay | Normal Medium | **6F** | 0.5 ± 0.1 |
| Alisertib Resistant +Alisertib | H2B-GFP decay | Normal Medium | **6F** | 0.6 ± 0.1 |
| Volasertib Resistant +Volasertib | H2B-GFP decay | Normal Medium | **6F** | 0.7 ± 0.1 |
| Ispinesib Resistant | H2B-GFP decay | Normal Medium | **6F** | 1.7 ± 0.1 |
| Alisertib Resistant | H2B-GFP decay | Normal Medium | **6F** | 3.6 ± 0.3 |
| Volasertib Resistant | H2B-GFP decay | Normal Medium | **6F** | 2.6 ± 0.2 |
| Ispinesib Resistant | H2B-GFP decay | Normal Medium | **6G** | 1.7 ± 0.1 |
| Ispinesib Resistant + Saracatinib | H2B-GFP decay | Normal Medium | **6G** | 3.0 ± 0.4 |
| Ispinesib Resistant | H2B-GFP decay | Normal Medium | **6H** | 1.9 ± 0.1 |
| Ispinesib Resistant + SB431542 | H2B-GFP decay | Normal Medium | **6H** | 2.8 ± 0.2 |
| Alisertib Resistant | H2B-GFP decay | Normal Medium | **6H** | 4.6 ± 0.2 |
| Alisertib Resistant +SB431542 | H2B-GFP decay | Normal Medium | **6H** | 5.8 ± 0.3 |
| Volasertib Resistant | H2B-GFP decay | Normal Medium | **6H** | 3.5 ± 0.2 |
| Volasertib Resistant + SB431542 | H2B-GFP decay | Normal Medium | **6H** | 5.1 ± 0.3 |

### Table S3: Rate Constants for L1 Cells

| Status | Process | Culture Condition | Fig. | Rate ($hr^{-1}$) x $10^3$ (± SEM) |
|---|---|---|---|---|
| Naive | H2B-GFP decay | Normal Medium | **S6B** | 5.3 ± 0.1 |
| Ispinesib Resistant + Ispinesib | H2B-GFP decay | Normal Medium | | 0.09 ± 0.01 |
| Alisertib Resistant + Alisertib | H2B-GFP decay | Normal Medium | | 0.06 ± 0.01 |
| Volasertib Resistant + Volasertibt | H2B-GFP decay | Normal Medium | | 0.08 ± 0.01 |
| Ispinesib Resistant | H2B-GFP decay | Normal Medium | **S6B** | 1.5 ± 0.1 |
| Alisertib Resistant | H2B-GFP decay | Normal Medium | **S6B** | 1.0 ± 0.1 |
| Volasertib Resistant | H2B-GFP decay | Normal Medium | **S6B** | 1.9 ± 0.1 |
| Ispinesib Resistant + SB431542 | H2B-GFP decay | Normal Medium | **S6C** | 2.2 ± 0.2 |
| Alisertib Resistant + SB431542 | H2B-GFP decay | Normal Medium | **S6C** | 1.9 ± 0.2 |
| Volasertib Resistant +SB431542 | H2B-GFP decay | Normal Medium | **S6C** | 2.7± 0.2 |

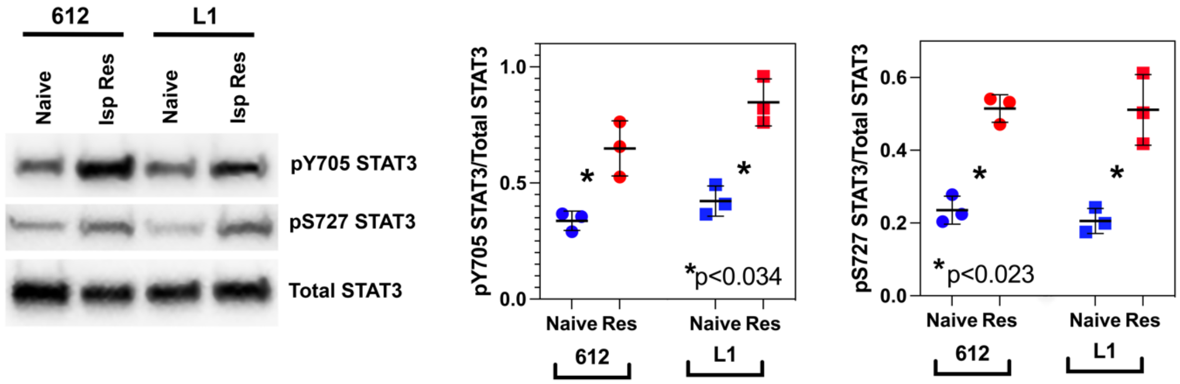

**Fig. S1: Western blots for pY705 STAT3, pS727 STAT3, and total STAT3 in human 612 and L1 GBM cell lines.** For both lines, ispinesib resistance is associated with a statistically significant increase in both phosphorylated species, compared to drug naïve cells.

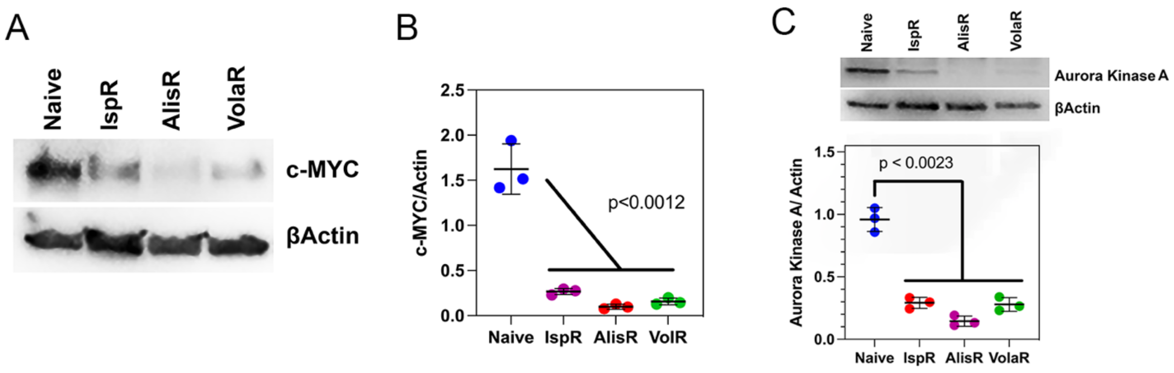

**Fig. S2:** Resistance to spindle inhibitors downregulates c-MYC by 8-10-fold (**A,B**) and downregulates expression of Aurora Kinase A 3-4 fold (**C**).

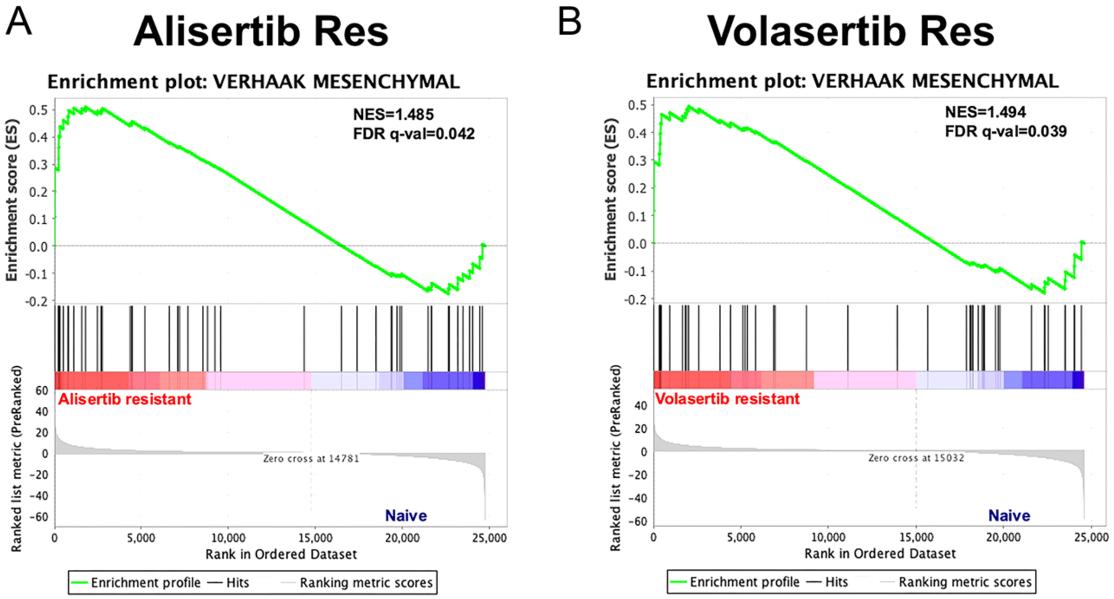

**Figure S3:** Gene set enrichment analysis of alisertib (A) and volasertib (B) resistant *Trp53/Pten*(-/-) GBM cells. Although the *Trp53/Pten*(-/-) murine GBM line has a strong proneural transcriptional signature, resistance to these two spindle inhibitors, as in the case of ispinesib (Kenchappa *et al.,* 2022) produces a proneural→mesenchymal transcriptional shift.

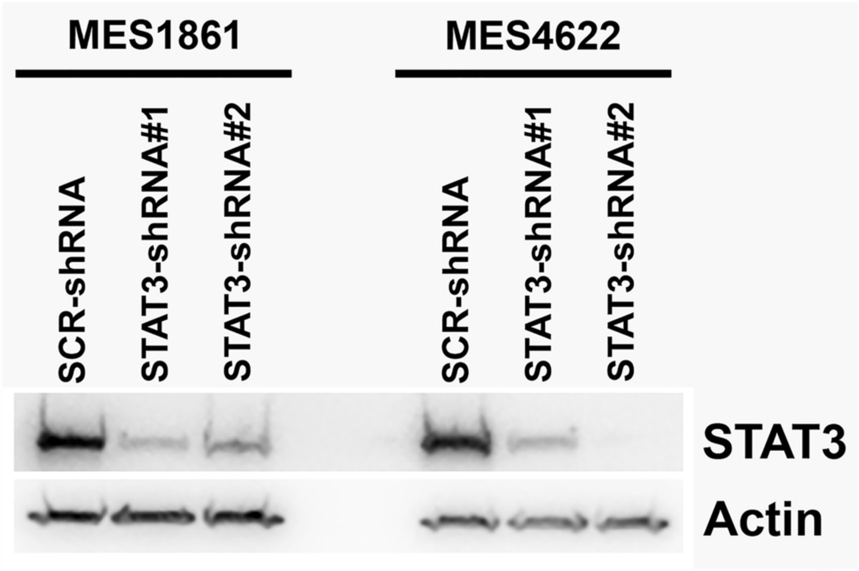

**Figure S4:** Western blots of STAT3 confirm that both targeting shRNAs reduce STAT3 levels by >90%.

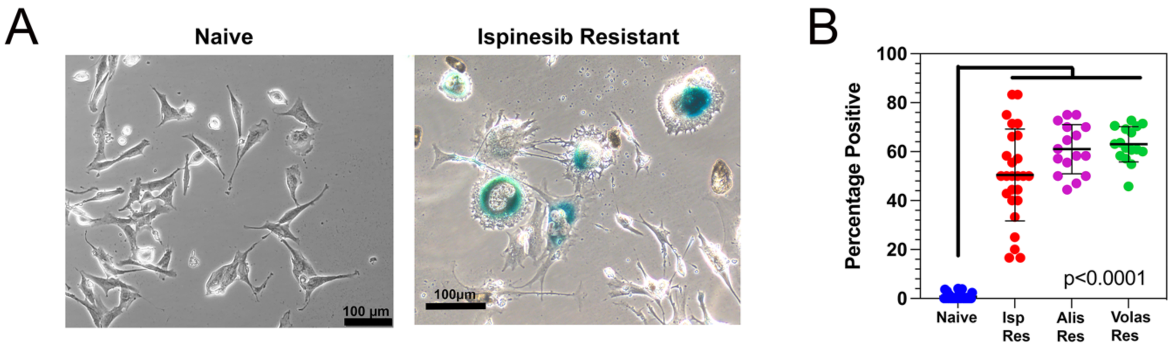

**Figure S5:** (**A**). X-Gal staining of drug naïve (*left*) and ispinesib resistant (*right*) *Trp53/Pten*(-/-) cells. (**B**). Quantitation of β-gal positivity demonstrates 55-65% positivity in ispinesib (*red*), alisertib (*magenta*), and volasertib (*green*) resistant cells.

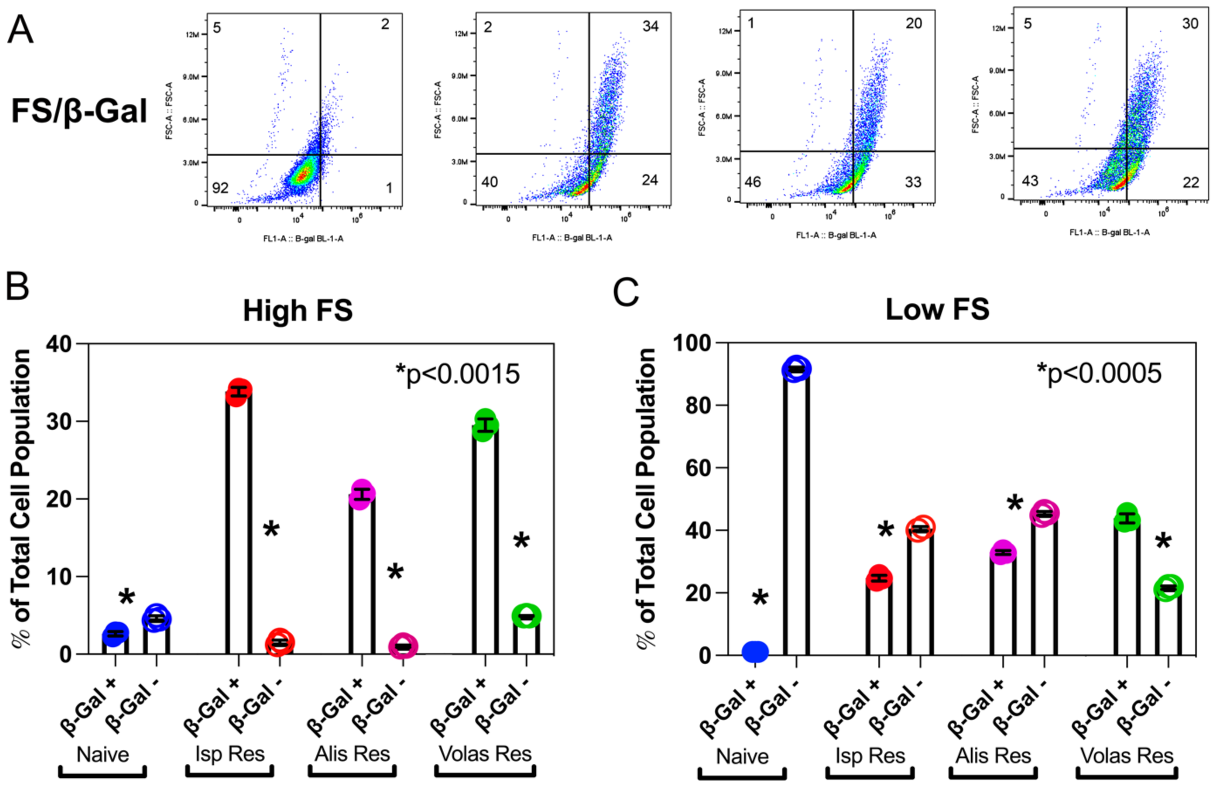

**Figure S6:** (**A**). Naïve and spindle inhibitor resistant *Trp53/Pten*(-/-) cells were stained with the fluorescent β-gal substrate FDGlu and subjected to flow cytometry to measure forward scatter *versus* fluorescence. A substantial fraction of the resistant cells demonstrate both high forward scatter and β-gal activity. Numbers in each quadrant represent the mean percentages of the total signal. (**B**). Bar plot depicting for naïve and resistant cells the fraction of the total cell population that demonstrates both high forward scatter and either positive or negative FDGlu fluorescence. (**C**). Bar plot depicting for naïve and resistant cells the fraction of the total cell population that demonstrates both low forward scatter and either positive or negative FDGlu fluorescence.

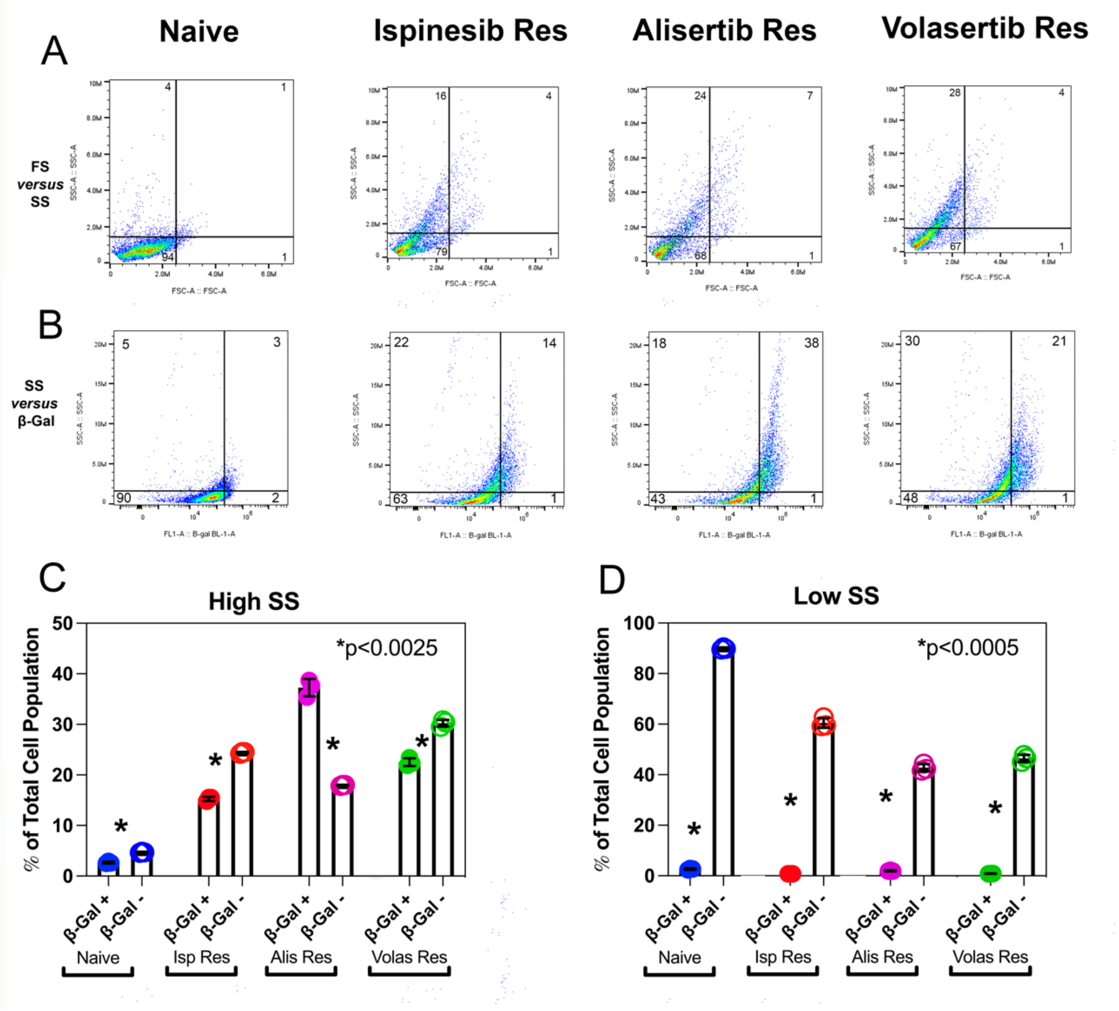

**Figure S7**: (**A**). Forward versus side scatter flow cytometry of L1 human GBM cells shows that while drug naïve cells cluster in a region of low forward and side scatter, resistance to each of the three spindle inhibitors generates a broad distribution of cells that include a 4-5 fold increase in cells with a high side scatter profile. Numbers in each quadrant represent the mean percentages of the total signal. (**B**). Naïve and resistant cells were stained with the fluorescent β-gal substrate FDGlu and subjected to flow cytometry to measure side scatter *versus* fluorescence. A substantial fraction of the resistant cells demonstrate both high side scatter and β-gal activity. Numbers in each quadrant represent the mean percentages of the total signal. (**C**). Bar plot depicting for naïve and resistant cells the fraction of the total cell population that demonstrates both high side scatter and either positive or negative FDGlu fluorescence. (**D**). Bar plot depicting for naïve and resistant cells the fraction of the total cell population that demonstrates both low side scatter and either positive or negative FDGlu fluorescence.

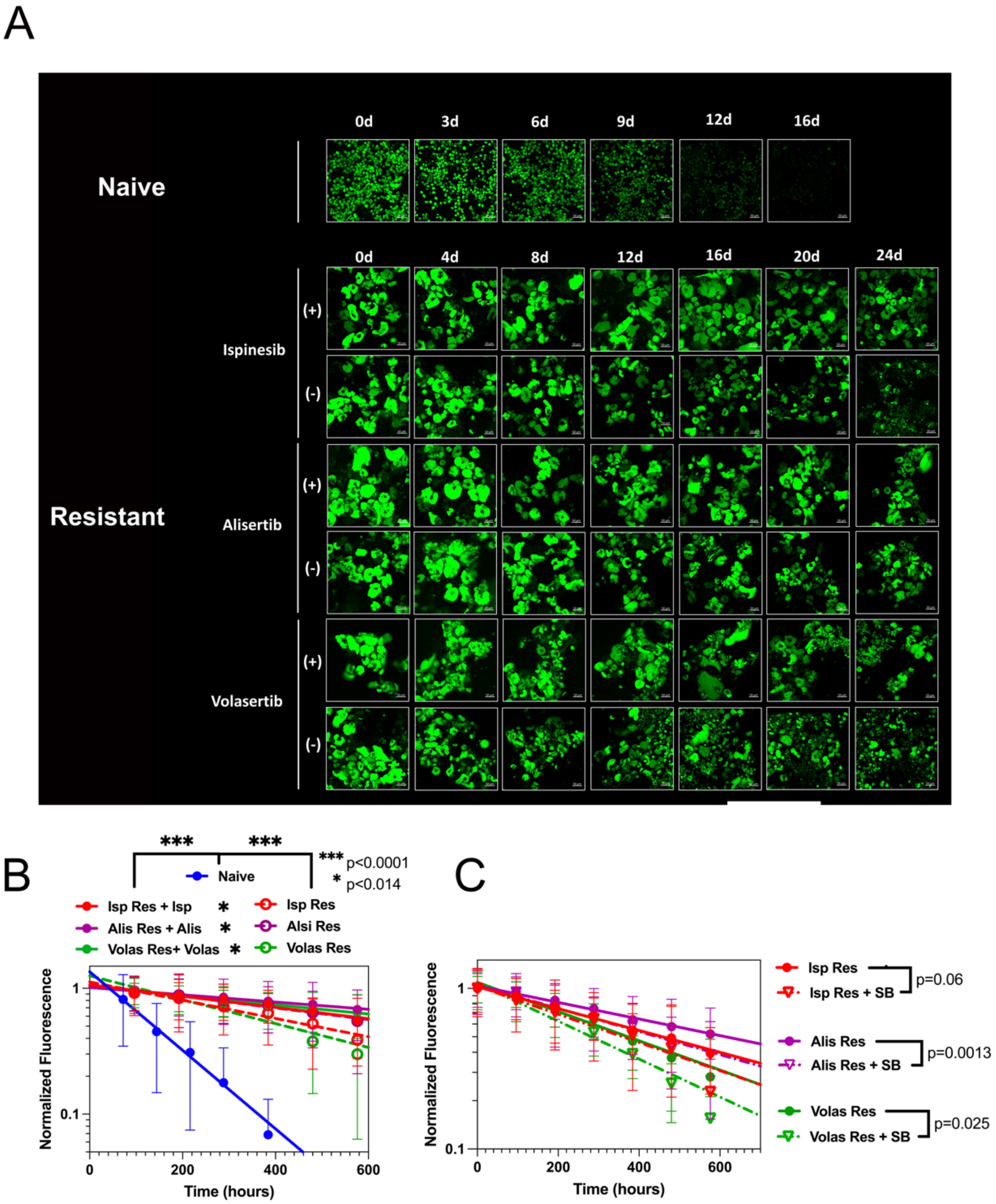

**Figure S8:** (**A**). Fluorescence micrographs of naïve and spindle inhibitor resistant human L1 GBM cells transfected with doxycycline inducible H2B-GFP. Cells were pulsed with doxycycline, and fluorescence intensity was measured after doxycycline removal, both in the presence (+) and absence (-) of spindle inhibitor. (**B**). Semi logarithmic plot of fluorescence decay of drug naïve (*blue*), ispinesib (red), alisertib (*magenta*), and volasertib (*green*) resistant L1 cells in the presence (*solid circles*) and absence (*open circles*) of spindle inhibitor. Corresponding rate constants from single exponential decays (*lines*) are summarized in **Table S3**. Proliferation of ispinesib (*red*), alisertib (*magenta*), and volasertib (*green*) resistant L1 cells in the absence of spindle inhibitor is accelerated 45-90% by addition of the TGFβ receptor inhibitor SB431542 (**Table S3**).
